# Mycobacteriophage Adephagia gp48 inhibits essential host cell wall biosynthesis enzyme mannose-1-phosphate guanylyltransferase

**DOI:** 10.64898/2026.09.16.752054

**Authors:** Michael J. Lauer, Krista G. Freeman, Danny Jiang, Jindanuch Maneekul, Andrew P. VanDemark, Graham F. Hatfull

**Affiliations:** Department of Biological Sciences, University of Pittsburgh, Pittsburgh PA 15260; Department of Physics, Case Western Reserve University, Cleveland OH 44106

## Abstract

It is common for bacteriophages to encode proteins that are strongly inhibitory to growth of the bacterial host, and about 10% of mycobacteriophage-encoded proteins have this property. Adephagia is a Cluster K1 mycobacteriophage and prior cloning and expression of 66 Adephagia non-structural genes identified 14 that are toxic when expressed in *Mycobacterium smegmatis*. One of the expressed proteins, the 70-residue gp48, is highly toxic when expressed in both *M. smegmatis* and *Mycobacterium abscessus* and acts by binding to and inactivating the function of Msmeg_1828, a mannose-1-phosphate guanylyltransferase (Mpg). Mpg is an essential enzyme for biosynthesis of GDP-mannose, a precursor of several key cell wall constituents including lipoarabinomannan and phosphatidylinositol mannosides. The crystal structure of Adephagia gp48 shows that the N-terminal 43 residues form two alpha helices that strongly promote dimer formation; the C-terminal 27 residues are disordered but are predicted to bind Fe-S clusters via cysteine and histidine residues. Adephagia gp48 inhibits Mpg guanylyltransferase activity *in vitro* and is predicted to interrupt Mpg dimer formation. The C-terminal metal binding activity of gp48 is not required for toxicity, and non-toxic mutants have substitutions in the N-terminal alpha helices that are involved in dimerization. The selective advantage of Mpg inactivation for the phage is unclear, but it may protect from competing phages that require GDP-mannose derived molecules for efficient infection.

**IMPORTANCE:** A majority of bacteriophage-encoded proteins are of unknown function. An effective strategy for gaining functional insights into these is to identify those that are toxic when expressed in the bacterial host and determine the basis of the toxicity. We show that gp48 encoded by mycobacteriophage Adephagia is toxic when overexpressed in *Mycobacterium smegmatis* and *Mycobacterium abscessus* and the toxicity results from gp48 interaction with the host mannose-1-phosphate guanylyltransferase (Mpg) enzyme. Mpg is required for biosynthesis of GDP-Mannose, a precursor for many essential mannose-containing cellular constituents, and gp48 acts a dimer to binding to Mgp and inactivate its enzymatic activity.

## INTRODUCTION

Mycobacterial infections are challenging to treat clinically, with inherent and acquired antibiotic resistance being common (1, 2). The isolation and characterization of mycobacteriophages have provided insights into these pathogens, produced tools for genetic analyses, and made possible new treatments for nontuberculous mycobacterium (NTM) infections (3–6). A large number of mycobacteriophages have been isolated – mostly using *Mycobacterium smegmatis* – and over 2,800 have been sequenced and annotated (7). They are genetically diverse and can be grouped into 36 clusters of closely related phages and five singletons, each with no close relatives (8). The majority of the cluster/singleton groups are temperate, and most have siphoviral morphologies. The virion structural protein genes are typically recognizable and organized into a late-expressed operon, and some non-structural genes such as integrases and DNA Polymerases can be readily identified. However, the genomes typically contain large numbers of genes of that are relatively small and are of unknown function (7).

One approach to gain insights into the functions of these phage proteins is to clone and overexpress individual proteins in a *Mycobacterium* host, typically *M. smegmatis* (9, 10). It is estimated that approximately 10% of all mycobacteriophage proteins exhibit toxicity when overexpressed in the host cell (10). Whole genome screens and other studies have identified many mycobacteriophage genes that are toxic when inducibly expressed in *M. smegmatis* (10–19). The reason why phages encode such genes is unclear – because the host bacterium will be dead once lysis occurs – but there are several examples where toxicity results from binding to and inactivating essential host functions. In the example of phage Fruitloop, the protein gp52 binds specifically to host protein DivIVA, and excludes infection by an unrelated phage, Rosebush, that requires DivIVA for infection (19). It is plausible that many cytotoxic proteins act similarly to prevent other phages from infecting and stealing valuable resources while the phage is lytically propagating. However, there may be considerable specificity for particular phages, making it challenging to identify these phage-phage dynamics. We note also that proteins that don’t exhibit such toxicity could behave similarly, interfering with host functions that are required for phage infection but which are not essential for viability.

Adephagia is a temperate Cluster K1 phage isolated on *M. smegmatis*, which also infects *M. tuberculosis* and some strains of *M. abscessus* (7, 20, 21). It has a 59,646 bp genome with 94 protein-coding genes and one tRNA gene (20). A screen for cytotoxic genes identified fourteen with toxic phenotypes, although the function of most of these has not be elucidated (17). Here, we explore the role of one of these cytotoxic proteins, the 70-residue Adephagia gp48, using bioinformatic, biochemical, structural, molecular, and genetic approaches to determine its function. Adephagia gene *48* could be readily deleted from the genome and we observed no interruption to lytic growth or the establishment of lysogeny. Immunoprecipitation of His-tagged gp48 from *M. smegmatis* pulled down a single additional protein identified as Msmeg_1828, a mannose-1-phosphate guanylyltransferase (Mpg). Simple addition of purified gp48 to purified Mpg showed that it not only binds but that it interrupts the enzyme activity of Mpg; we also isolated non-toxic gp48 mutants that bind but preserve enzyme activity. AlphaFold3 predicts that a gp48 dimer inhibits Mpg dimerization, and the gp48 non-toxic mutants map primarily in the gp48 dimerization interface, suggesting the importance of dimerization in the protein’s mechanism of action.

## RESULTS

### Adephagia gp48 expression is cytotoxic to *M. smegmatis* and *M. abscessus* GD40

One of the Adephagia genes shown previously to be toxic when expressed in *M. smegmatis* is *48* (17), a 213-bp ORF coding for a predicted 70-residue protein; its function is unknown, but it is the fifth gene in the early lytic operon and is expressed early in lytic growth (17). We confirmed its toxic phenotype by transforming plasmid pKF175 – in which *48* is ATc-inducible – into *M. smegmatis* and showed that induction of gp48 expression is strongly growth inhibitory (Fig. 1A). When gp48 expression is induced in liquid culture, bacterial growth ceases after a short period of additional growth (Fig. 1A). We also transformed pKF175 into a clinical isolate of *M. abscessus* (GD40) and showed that induction also results in loss of viability, although the inhibition is milder than in *M. smegmatis*, both in recovery on solid media and in liquid culture (Fig. 1B). To determine if this phenotype is bactericidal or bacteriostatic, *M. smegmatis* mc^2^155 cells carrying pKF175 were induced in a liquid medium and aliquots plated on solid media with and without ATc at different times after induction (Fig. 1C). Substantial recovery was observed on solid media without ATc even after eight hours of induction in liquid culture suggesting that the toxicity is largely bacteriostatic (Fig. 1C).

**Figure 1.**
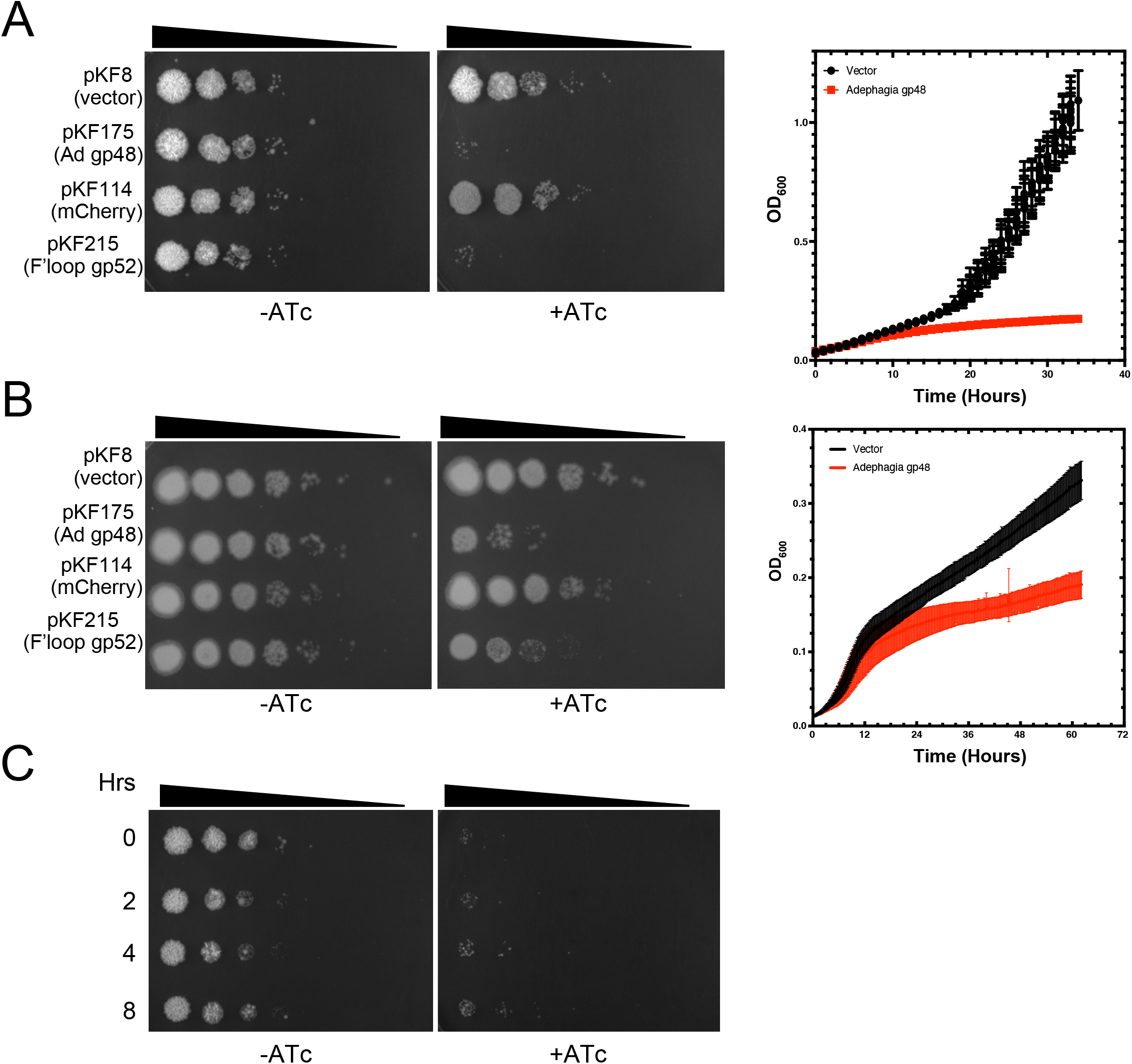
Cytotoxicity of Adephagia gp48. **A.** Ten-fold serial dilutions of cultures of *M. smegmatis* carrying plasmids (as indicated on the left) were plated on solid media with or without the ATc inducer as shown. On the right is shown growth curves in liquid medium with ATc of three independent cultures each of colonies carrying either the empty vector (black) or ATc-inducible gp48 (red). Data are shown as average points with standard deviation error bars **B.** Ten-fold serial dilutions of cultures of *M. abscessus* GD40 carrying plasmids (as indicated on the left) were plated on solid media with or without the ATc inducer as shown. On the right is shown growth curves in liquid medium with ATc of three independent cultures each of colonies carrying either the empty vector (black) or ATc-inducible gp48 (red). **C.** Adephagia gp48 expression is mostly bacteriostatic in *M. smegmatis* mc^2^155. Liquid cultures of *M. smegmatis* carrying plasmid pKF175 with the ATc inducer were sampled at varying times after induction (as shown at the left), ten-folds serially diluted, and plated onto solid media with without ATc inducer (left) or with ATc inducer (right).

### Mycobacteriophage homologues of Adephagia gp48

Grouping of actinobacteriophage proteins into phamilies (phams) according to sequence similarity (22, 23) places Adephagia gp48 into a phamily (current designated Pham 850) with 143 other proteins. The related proteins are predominantly in Cluster K phages (20), specifically within Subclusters K1, K2, and K6, but not in Subclusters K3, K4, K5, K7 or K8 (Fig. 2A). Nearly all the K1 and K2 phages carry a homologue, but only about 50% of the 26 Subcluster K6 phages do. There is also a phamily member in phage Nergal, one of two phages in Cluster AG (Fig. 2A). In all instances, the gene is located 4-5 genes downstream from the start of the putative rightwards early lytic promoter (Fig. 2A). Alignment of the related sequences and the phylogenetic relationships (Figs. 2B, C, S1, S2) shows them to be quite diverse in sequence, with small insertions in some members relative to the others (Figs. 2B, C, S1, S2). We note that the Hammy gp51 and Amelie gp44 homologues are also cytotoxic when overexpressed in *M. smegmatis* mc^2^155 (14, 16) – other phamily members remain to be tested.

**Figure 2.**
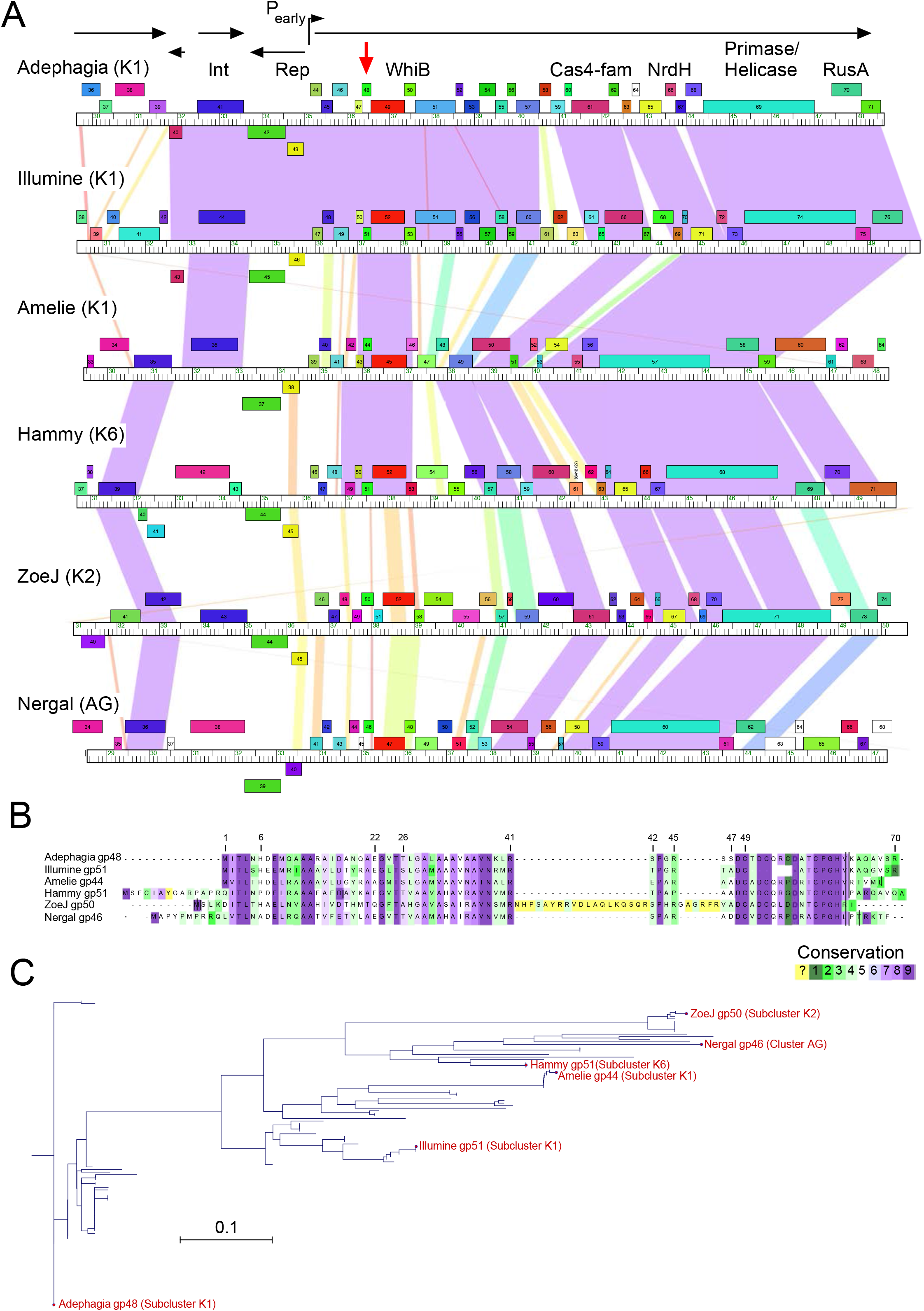
Adephagia gp48 conservation within Cluster K mycobacteriophages. **A**. Partial genome organizations of phages Adephagia, Illumine, Amelie, Hammy, ZoeJ, and Nergal; Cluster and subcluster designations are shown after phage names on the left. Genes are shown as colored boxes above and below each genome ruler, reflecting rightwards and leftwards transcription, respectively, and designated by arrows with features such as the promoter P_early_. Pairwise BLASTN nucleotide similarity is shown as spectrum colored shading between genomes, with violet being the most similar and red the least similar above a threshold E-value of 10^-4^. Gene functions, where known, are indicated above genes. Adephagia gene *48* is indicated with a red vertical arrow. **B**. Amino acid sequence alignments of Adephagia gp48 and related proteins colored by conservation among all 144 pham 850 members using a ConSurf evolutionary conservation mapping for each amino acid, scaled 1-9; 1 is least conserved (dark green), 5 is variable (white), 9 is most conserved (purple), and yellow is variable. **C**. Phylogenetic tree constructed from primary amino acid sequence alignment of 144 members of pham 850 using maximum likelihood method with Jukes-Cantor protein distance measure and bootstrap analysis of 500 replicates, showing 6 representative proteins: Adephagia gp48, Illumine gp51, Amelie gp44, Hammy gp51, ZoeJ gp50, and Nergal gp46.

### Adephagia gp48 is not required for lytic growth or lysogeny

To determine if gene *48* is required for Adephagia lytic growth we constructed a derivative (AdephagiaΔ*48*) in which the gene is deleted (Fig. 3). Using bacteriophage recombineering of electroporated DNA (BRED) (24) we co-electroporated a 500bp synthetic DNA substrate containing the Δ*48* deletion and flanking sequences (Fig. 3A) with Adephagia genomic DNA, then plated on an *M. smegmatis* lawn to recover primary plaques. Six of these were screened by PCR and a plaque containing the mutant allele was re-purified and secondary plaques tested by PCR (Fig. 3B). A purified lysate was sequenced to confirm the genome was as predicted. We repeated the construction using genomic DNA of an AdephagiaΔ*43* lytic derivative (AdΔ*43*) to make Adephagia Δ*43*Δ*48* (Fig. 3B). The constructions of the Δ*48* derivatives were straightforward, strongly suggesting that gene *48* is not required for lytic growth. Mutants and their wild type cognate parents form plaques with similar appearance, and the plaques contain similar numbers of particles, although the Δ48 plaques are somewhat more variable in their particle numbers (Fig. 3C).

**Figure 3.**
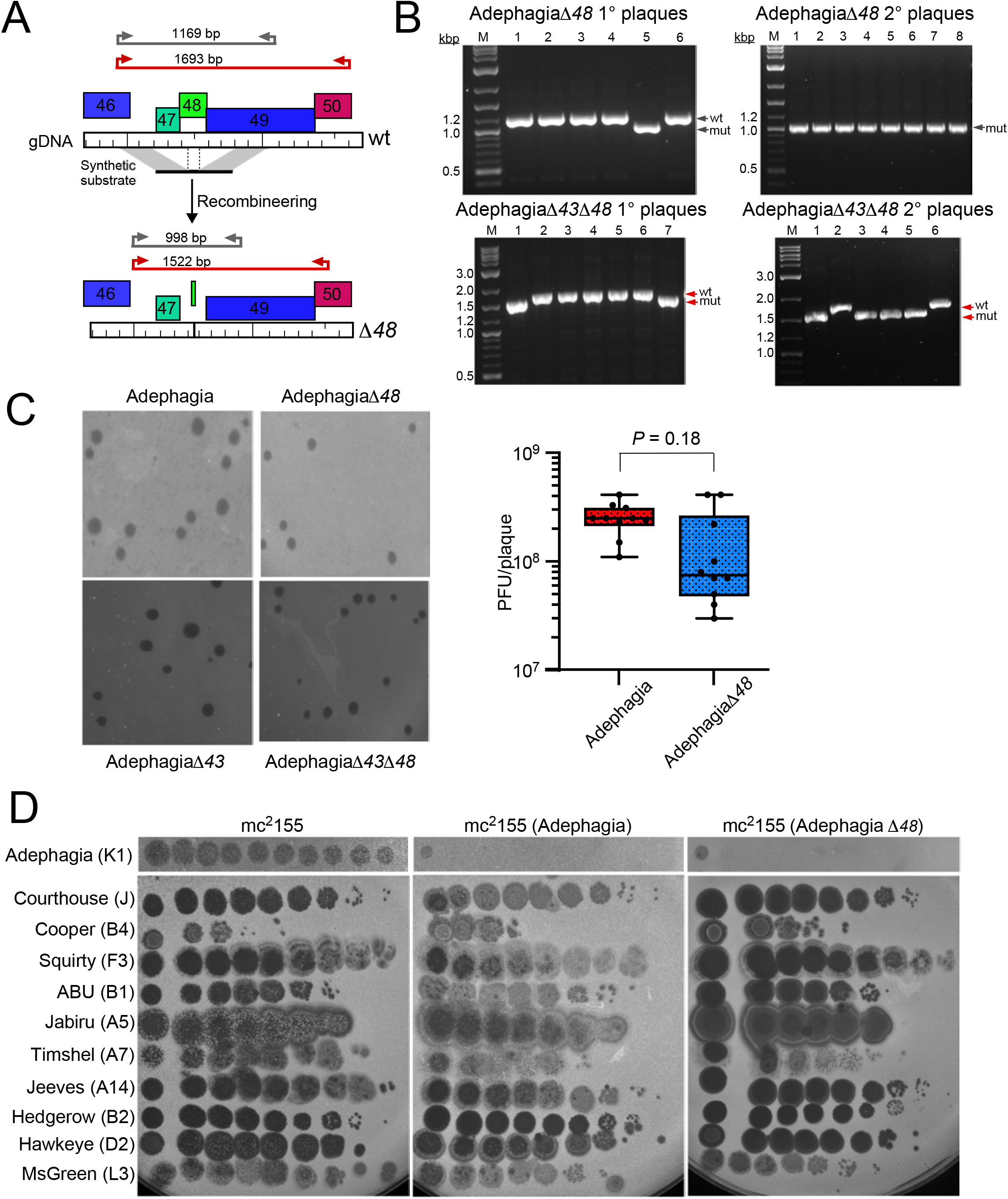
Adephagia gp48 is not required for lytic growth or lysogeny. **A.** Schematic illustrating the construction of Adephagia gene *48* deletion mutants by Bacteriophage Recombineering of Electroporated DNA (BRED). The top panel depicts a segment of the Adephagia genome around *48*, with two primer sets shown in grey and red, with gray shading indicating the homologous regions included in the synthetic substrate. The bottom panel shows the Δ*48* mutant genomes with *48* deleted and the same primer sets denoted with new amplicon sizes. **B**. PCR screening of primary (left) and secondary (right) plaques recovered and isolated from the temperate Δ*48* (top) and the lytic Δ*43*Δ*48* (bottom) BRED reactions, with wildtype and mutant bands indicated with arrows colored corresponding to primer sets in panel A. **C.** Plaque phenotypes of Adephagia, including isolated recombinant deletion mutants. Temperate phages are shown on the top row (wildtype, left and Δ*48*, right) and lytic phages are shown on the bottom row (Δ*43*, left and Δ*43*Δ*48*, right). To the right, the number of plaque forming units released from resuspended single plaques of lytic AdephagiaΔ*43* (wildtype) or AdephagiaΔ*43*Δ*48* (deletion mutant) phages (n=10) are shown. **D.** Ten-fold serial dilutions of Adephagia and a panel of 10 unrelated phages with their cluster and subclusters denoted spotted onto lawns of *M. smegmatis* mc^2^155 (left), or lysogens of Adephagia: wildtype (middle) or Adephagia Δ*48* (right).

We could readily isolate lysogenic strains carrying either Adephagia wild type or AdephagiaΔ*48* prophages and gp48 is clearly not required to establish lysogeny (Fig. 3D). The lysogens appear to grow normally and confer immunity to Adephagia superinfection, such that gp48 is not involved in superinfection immunity or homotypic exclusion (Fig. 3D). We also compared the lysogens for superinfection by a diverse suite of other mycobacteriophages (Figs. 3D, S3) to ask if gp48 is involved in prophage-mediated phage defense. We observed no differences and no evidence for such a role.

### Adephagia gp48 is cytoplasmic and interacts with Msmeg_1828 (Mpg; ManB)

To further explore the toxicity of Adephagia gp48, we constructed an N-terminal protein fusion with mCherry expressed from an ATc-inducible promoter that retains its toxicity (Fig. S4). On induction of Adephagia gp48 expression in liquid growth we saw no major changes to cell shape or morphology, and fluorescence was broadly distributed throughout the cytoplasm (Fig. 4A). Although the fluorescence occasionally has a peapod-like appearance we saw no distinct puncta or membrane localization (Fig. 4A).

**Figure 4.**
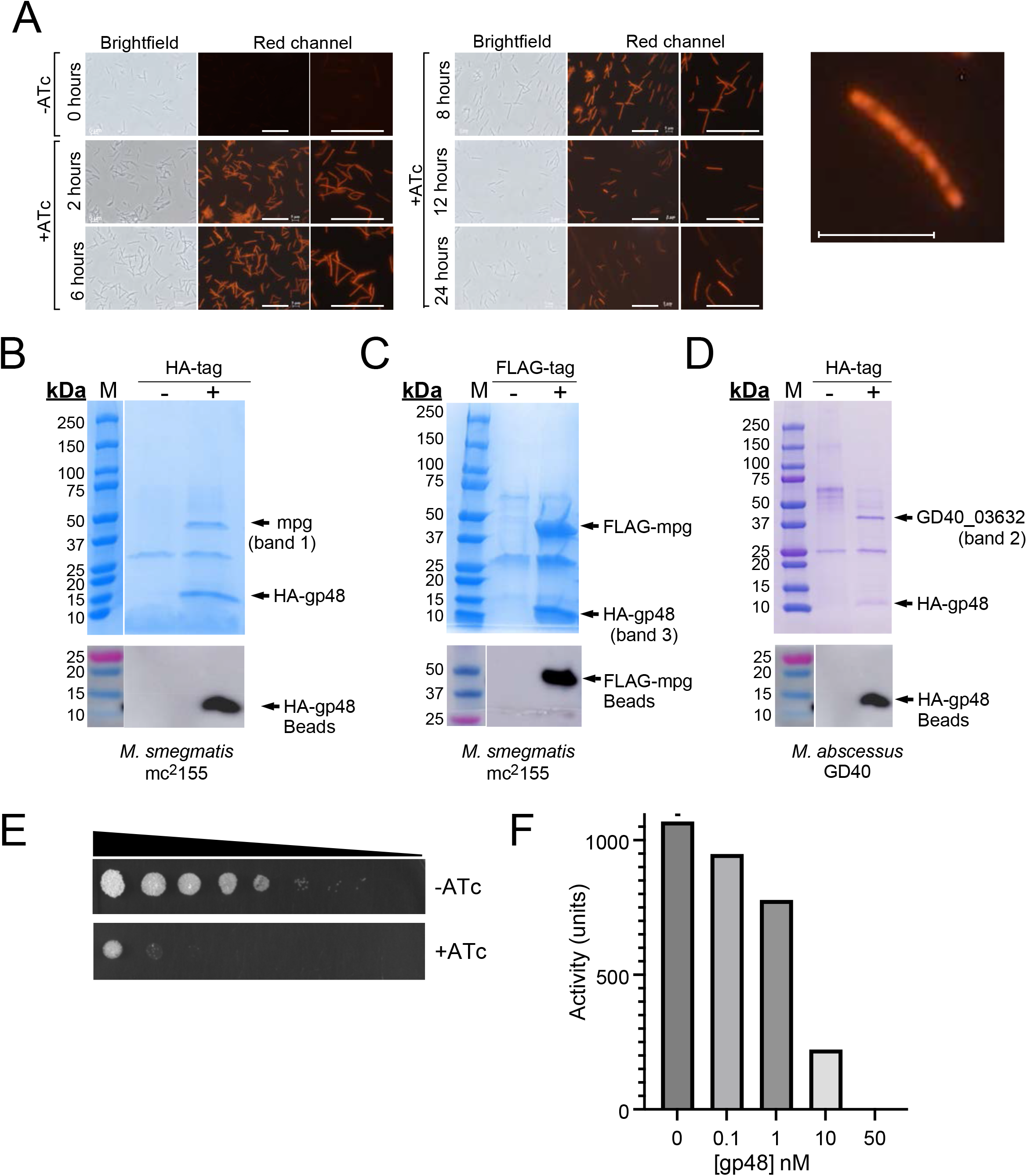
Adephagia gp48 is cytoplasmic and interacts with Mpg. **A.** Cells expressing an N-terminal-tagged mCherry Adephagia gp48 were induced and cells were mounted and imaged at time 0 (uninduced), 2, 6, 8, 12, and 24 hours post-induction on brightfield channel (left) and fluorescence channels for mCherry at two different magnifications. Scale markers are 20 µm. Shown to the right is a higher magnification of the 24-hour sample, displaying a peapod appearance; scale bar is 5 µm. **B.** Clarified lysates were prepared from *M. smegmatis*, expressing either untagged Adephagia gp48 (-) or Adephagia gp48 with an N-terminal HA tag (+). Following immunoprecipitation and washing, proteins were separated on a 4-20% SDS-PAGE gel and stained with Imperial Blue. An arrow indicates the expected migration of Adephagia HA-gp48, and a second arrow shows a prominent band (labeled band 1) absent in the non-tagged control that was identified by LC/MS-MS as Msmeg_1828 (mpg) in *M. smegmatis*. Shown below is a Western blot of the HA-gp48 pulldown using anti-HA beads from these eluate lanes. **C.** Anti-FLAG pulldown with untagged Msmeg_1828, or FLAG-tagged Msmeg_1828 in *M. smegmatis*, each co-expressing HA-tagged Adephagia gp48 (band 3 sent for LC-MS/MS identification) and a Western blot of anti-FLAG bead eluates from the FLAG-mpg reverse pulldown, confirming the identity of the bait protein. **D.** Eluates of anti-HA beads from similarly prepared *M. abscessus* GD40 cell lysates expressing either untagged Adephagia gp48 or HA-gp48 featuring an arrow that indicates the expected migration of Adephagia HA-gp48, and a second arrow that shows a prominent band absent in the non-tagged control. That band (labeled band 2) was identified by LC/MS-MS as GD40_03632 in *M. abscessus* GD40, a homolog of mannose-1-phosphate guanylyltransferase. Presence of HA-gp48 bait is demonstrated below by Western blot of the HA-bead eluate lanes. **E.** Ten-fold serial dilutions of *M. smegmatis* cultures expressing a CRISPRi knockdown system targeting Msmeg_1828 on solid media with (+) or without (-) inducer ATc. **F.** Purified Msmeg_1828 protein and substrates (500 µM mannose-1-phosphate, 250 µM GTP, 2 mM MgCl_2_) were mixed with either varying concentrations of Adephagia gp48 (0.1, 1, 10, and 50 nM) or 750 nM BSA (0 nM gp48) as a control at 25°C and activity units λ_ex_ = 316 nm/λ_em_ = 456 nm were quantified 10 minutes post-mixture.

To determine if Adephagia gp48 interacts with *M. smegmatis* proteins, we expressed gp48 with an HA-tag at its N-terminus in *M. smegmatis* mc^2^155. We immunoprecipitated the fusion protein, collected associated proteins, and separated them by SDS-PAGE (Fig. 4B). In addition to gp48, we observed a prominent ∼40kDa protein, which is absent in a control lacking the HA tag (Fig. 4B). This protein was analyzed by LC-MS/MS and identified as Msmeg_1828 (Table 1). Msmeg_1828 was previously annotated as a pseudogene in *M. smegmatis* but was subsequently shown to be a 359-residue protein with a 37.5 kDa predicted mass (25). The gene codes for a mannose-1-phosphate guanylyltransferase (Mpg) enzyme involved in forming GDP-mannose that is required for synthesis of many cell wall components including phosphatidylinositol mannosides (PIMs), polyprenyl-phosphate-mannose (PPMs), lipomannan (LM), lipoarabinomannan (LAM), and methylmannose polysaccharides (MMPs) (26, 27).

**Table 1:** LC-MS/MS identification of Adephagia gp48 host binding partner.

| Gel band #<sup>1,2</sup> | Identified protein | Accession / locus | MW (Da) | Protein prob. | Unique peptides | Total Spectra | Sequence coverage |
| --- | --- | --- | --- | --- | --- | --- | --- |
| Band 1: *M. smegmatis* mc<sup>2</sup>155 (HA-gp48 IP) | Msmeg_1828 (Mpg) | AFP38256.1,YP_006566551.1 | 39,147 | 100% | 22 | 145 | 61.7% |
| Band 2: *M. abscessus* GD40 (HA-gp48 IP) | GD40_03632 (Mpg homolog) | GD40_03632 | 37,685 | 100% | 70 | 2258 | 99.7% |
| Band 3: *M. smegmatis* mc<sup>2</sup>155 (FLAG-Msmeg_1828 rev-IP) | HA-Adephagia gp48 | HA-Adephagia gp48 fusion | 9,744.6 | 100% | 4 | 22 | 43.0% |

<sup>1</sup>Co-immunoprecipitating bands were excised from the bait-epitope-tagged (+) lanes shown in Figure 4B-D and bait identity was confirmed by Western blot (Fig. 4, S5).
<sup>2</sup>Band 3 corresponds to the reverse co-immunoprecipitation in which FLAG-tagged Msmeg\_1828, not HA-gp48, was expressed as the bait (Fig. 4C); HA-gp48 was identified by LC-MS/MS and Msmeg\_1828 identity was confirmed separately by Western blot.

To confirm the Adephagia gp48-Mpg interaction, we constructed an N-terminally FLAG tagged version of Mpg and co-electroporated it with the HA-gp48 into *M. smegmatis,* precipitated the FLAG-Mpg fusion protein, and examined the associated proteins by SDS-PAGE (Fig. 4C). We observed both the FLAG-Mpg and HA-gp48 proteins (Fig. 4C), which were identified by Western blotting (Figs. 4, S5); the ∼10-kDa band was further confirmed by LC-MS/MS as Adephagia gp48 (Table 1). We repeated the immunoprecipitation in *M. abscessus* GD40 (Fig. 4D) and showed that the *M. abscessus* Mpg homologue (GD40_03632) co-precipitates with gp48 (Table 1). Given its central metabolic role, Mpg is anticipated to be essential for mycobacterial growth, and this has been shown for the *M. tuberculosis* homologue (Rv3264c, ManB) (28). We constructed a sgRNA-expressing CRISPRi plasmid targeting *Msmeg_1828* and showed that its expression is toxic in *M. smegmatis* (Fig. 4E). The toxicity of Adephagia gp48 can thus be directly attributed to its interaction with and inactivation of *M. smegmatis* Mpg.

### Adephagia gp48 inhibits Mpg activity *in vitro*

*Msmeg_1828* was cloned, expressed, and purified from *E. coli* to near homogeneity (Fig. S7), similar to previous reports (29, 30). When the *E. coli* purified Mpg protein is incubated with GTP and mannose-1-phosphate we observed time-dependent release of pyrophosphate (PPi), indicating that the protein is enzymatically active (Fig. 4F). When Adephagia gp48 was added to the reaction, we saw a robust reduction in activity (Fig. 4F), such that Adephagia gp48 toxicity results from a direct interaction with Mpg and interference with its essential activity of producing GDP-mannose.

### Crystal structure of Adephagia gp48

To characterize the Adephagia gp48 structure, we overexpressed a fusion protein with 10xHis-mRuby2 at its N-terminus fused to gp48 *via* a linker with a TEV cleavage site (Figs. 5, S6). The fusion protein expressed well in *E. coli* and was partially purified with Ni-NTA affinity chromatography and cleaved with TEV protease (Fig. 5A). The 10xHis-mRuby2 was removed with Ni-NTA affinity chromatography and gp48 was further purified using anion exchange chromatography (Fig. 5B). During the purification we observed that the protein elutes from an anion exchange column with the prominent species migrating with an apparent molecular mass of ∼11 kDa, slightly larger than its predicted mass of 7.2 kDa (Fig. 5B). Size exclusion chromatography showed the protein to elute with a major peak between 14.3 kDa and 27 kDa on a calibrated column (Fig. 5C), consistent with the protein being primarily dimeric in solution. The fractions collected from both anion exchange (Fig. 5B) and size exchange (Fig. 5C) columns contained small amounts of a slower migrating protein on the SDS-PAGE analyses, with an apparent molecular weight of ∼18 kDa, which may correspond to protein dimers not fully denatured by SDS. Additionally, the peak fractions from the chromatography (Figs. 5B, C) were noticeably golden brown in color, consistent with gp48 containing a metal. We note that the extreme C-terminal part of the protein contains four cysteines (C49, C52, C55, C59) and a nearby histidine residue (H62) some of which could participate in metal-binding (Fig. S6), although C55 is not conserved in related proteins (Figs. 2B, S2).

**Figure 5.**
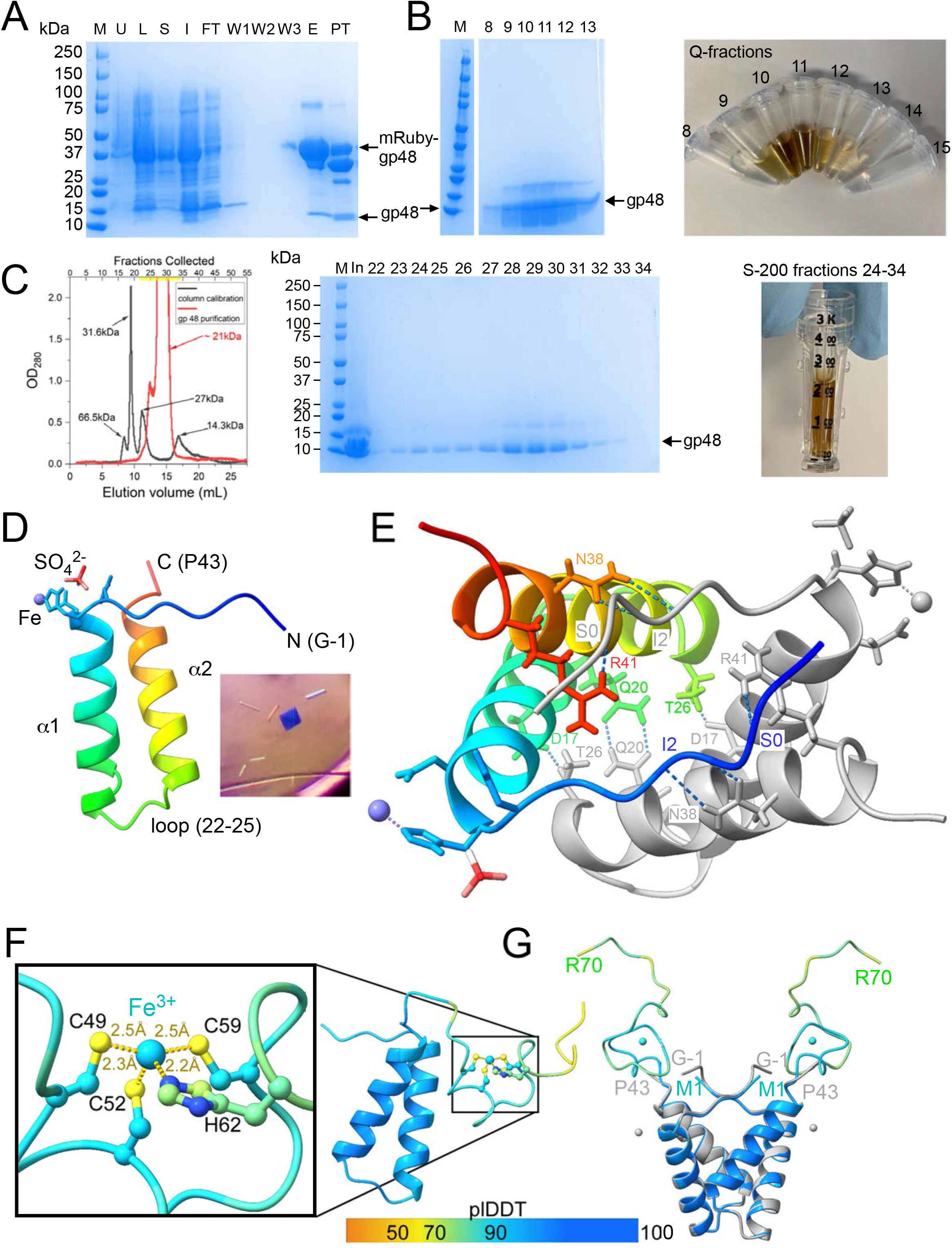
Purification and crystal structure of Adephagia gp48. Denatured gel samples were prepared from various stages of Adephagia gp48 affinity and Q ion-exchange chromatography purification and run on a 4-20% SDS-PAGE gel. **A.** The top left gel displays protein samples removed from uninduced (U) and autoinduced Rosetta DE3 cells expressing 10X-His-mRuby2-Adephagia gp48 fusion protein from cell lysate (L), soluble and insoluble lysate fractions (S, I), Ni-NTA column flowthrough (FT), NaCl buffer washes (W1, W2, W3), elution in 500 mM imidazole (E), and the elution cleaved with TEV protease (PT). **B.** The SDS-PAGE gel shows peak absorption and protein-containing fractions 8, 9, 10, 11, 12, and 13 from anion exchange chromatography over a steep 100-800mM NaCl gradient. The right panel demonstrates the hue of these fractions. **C.** A chromatogram shows the gp48 protein recovered from S-75 gel filtration, calibrated with four unrelated proteins (black line: lysozyme, 14.3 k-Da; TEV,27-kDa; Cytochrome c1, 31.6 kDa; BSA, 66.5 kDa). The gel to the right shows the pooled input to the S-75 column (In) and peak fractions as labeled. The concentrated peak fractions 22-34 are shown with a golden-brown hue to the right. **D.** The crystal structure of gp48 is shown as a monomer with bound sulfate and low-occupancy iron. The protein is shown as a cartoon model and colored blue (N-terminus, G-1) to red (C-terminus, P43); the two alpha helices are labeled (α1, α2), as is the loop connecting them (residues 22-25). An inset shows the protein crystals. **E.** The dimeric asymmetric unit is shown from the top down, with one subunit colored in rainbow as in panel D and the other in gray. The sidechains of residues involved in hydrogen bonds or close contacts that form the dimerization interface are shown as sticks, labeled, and connected with a dotted blue line. **F.** Predicted AlphaFold3 model of monomeric Adephagia gp48 and a single Fe^3+^ ion with intermolecular forces between C49, C52, C59, and H62 predicted to coordinate the ion shown in yellow dotted lines. The model is colored by prediction confidence indicated by pLDDT (as shown in key, with blue being high confidence). **G.** Predicted AlphaFold3 model of dimeric Adephagia gp48 with two Fe^3+^ ions colored by pLDDT (see key). The structure is aligned with the crystal structure of the Adephagia gp48 homodimer, shown in gray. The alpha helices of the predicted and experimentally determined structures are fully superimposable. The two Fe^3+^ ions in the Alphafold predicted structure are shown as blue spheres, and the two Fe^3+^ ions in the crystals are shown as grey spheres. The sulfur ions in the crystals are not shown.

Using purified gp48, we screened a diverse panel of crystallization conditions and observed square-shaped plate crystals after 21 days using a well solution containing 2 M ammonium sulfate, 100 mM Tris-HCl pH 7.0, and 200 mM lithium sulfate (Fig. 5D). X-ray diffraction data were collected on cryoprotected crystals, which belong to the space group C 1 2 1. Molecular replacement using portions of an AlphaFold3-predicted structure were used to estimate phases, and a structural model was refined to an R-work/R-free of 17.04% and 19.00% respectively. The asymmetric unit contains one copy of the gp48 protein, although only residues 1-43 of the 70-residue protein could be resolved, plus a Gly-Ser remaining after TEV cleavage (Table 2, Fig. S6). The C-terminal half of the protein (residues 44-70) was disordered and could not be modeled into electron density. The monomeric gp48 structure contains two alpha helices (α1,α2) connected by a short loop (residues 22-25) (Fig. 5E) and a coordinated ion, tentatively assigned as an iron ion (Fig. 5D). This ion is coordinated by His6-Nε2 and Asp7-Oδ1 (2.00 Å and 2.26 Å, respectively) and is situated in the plane of symmetry between two unit cells of the crystal, suggesting it could facilitate crystal packing. Analysis of packing within the gp48 crystal revealed a large interface between two gp48 subunits. This crystallographic interface buries 966 Å^2^ of surface area per protomer (1932 Å^2^ total) and contains more than half of the ordered residues, and in conjunction with gel filtration data, we conclude that gp48 is dimeric (Fig. 5E, Table S3). The dimerization interface is formed by polar contacts between side chains of opposing monomeric subunits, including hydrogen bonds: S0 and R41 (Ser0, positioned before Val1 is derived from the TEV cleavage site), I2 with N38, and Q20 with itself. Additional buried contacts, including D17 with T26 further stabilize the interface (Figs. 5E, S6, Table S3).

**Table 2:**
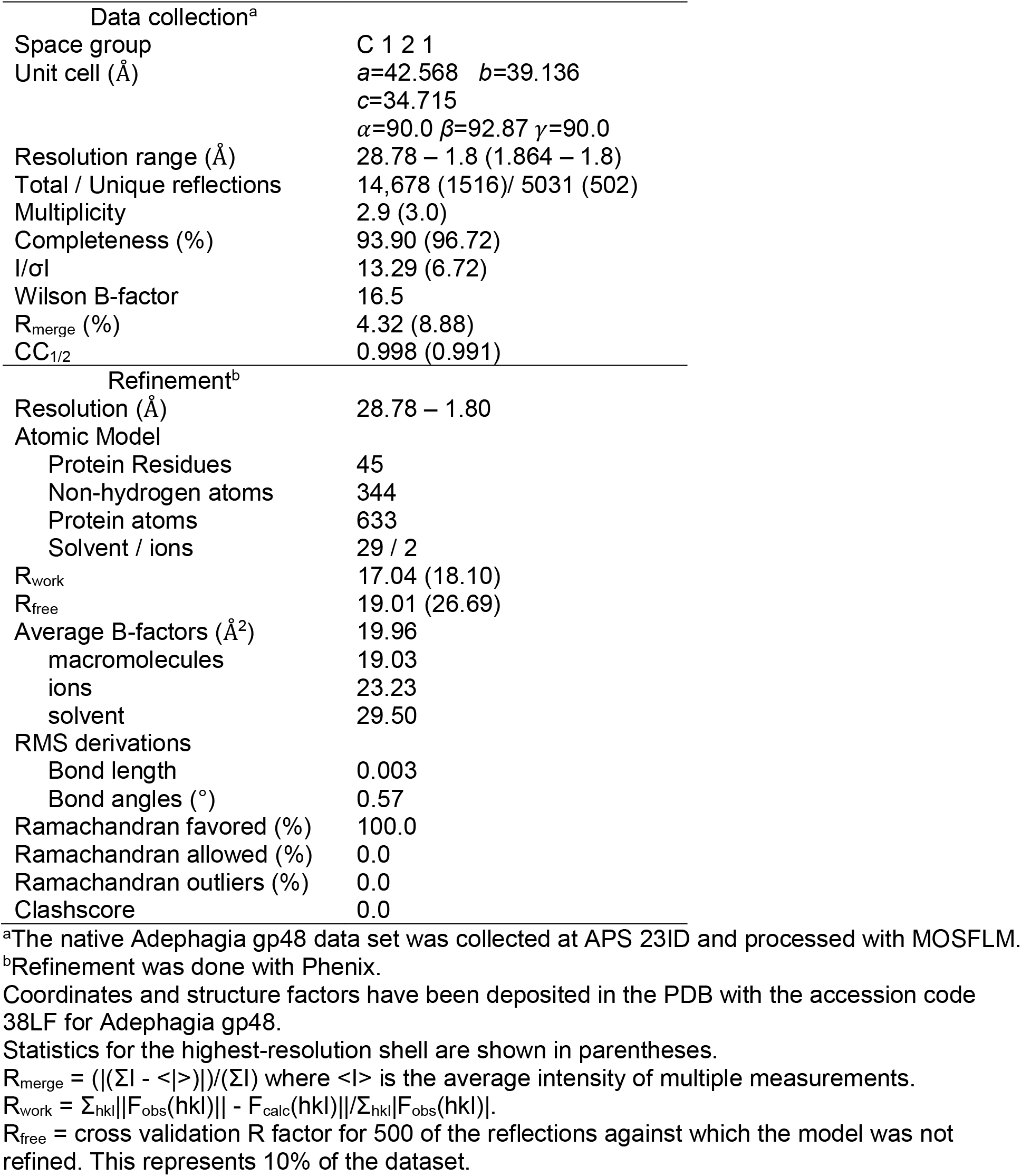
Adephagia gp48 data collection and refinement statistics.

| Data collection<sup>a</sup> |  |
| --- | --- |
| Space group | C 1 2 1 |
| Unit cell (Å) | *a*=42.568 *b*=39.136<br>*c*=34.715<br>$\alpha$=90.0 $\beta$=92.87 $\gamma$=90.0 |
| Resolution range (Å) | 28.78 – 1.8 (1.864 – 1.8) |
| Total / Unique reflections | 14,678 (1516)/ 5031 (502) |
| Multiplicity | 2.9 (3.0) |
| Completeness (%) | 93.90 (96.72) |
| I/σI | 13.29 (6.72) |
| Wilson B-factor | 16.5 |
| R<sub>merge</sub> (%) | 4.32 (8.88) |
| CC<sub>1/2</sub> | 0.998 (0.991) |
| Refinement<sup>b</sup> |  |
| Resolution (Å) | 28.78 – 1.80 |
| Atomic Model |  |
| Protein Residues | 45 |
| Non-hydrogen atoms | 344 |
| Protein atoms | 633 |
| Solvent / ions | 29 / 2 |
| R<sub>work</sub> | 17.04 (18.10) |
| R<sub>free</sub> | 19.01 (26.69) |
| Average B-factors (Å<sup>2</sup>) | 19.96 |
| macromolecules | 19.03 |
| ions | 23.23 |
| solvent | 29.50 |
| RMS derivations |  |
| Bond length | 0.003 |
| Bond angles (°) | 0.57 |
| Ramachandran favored (%) | 100.0 |
| Ramachandran allowed (%) | 0.0 |
| Ramachandran outliers (%) | 0.0 |
| Clashscore | 0.0 |

<sup>a</sup>The native Adephagia gp48 data set was collected at APS 23ID and processed with MOSFLM.
<sup>b</sup>Refinement was done with Phenix.
Coordinates and structure factors have been deposited in the PDB with the accession code 38LF for Adephagia gp48.
Statistics for the highest-resolution shell are shown in parentheses.
$R_{\text{merge}} = (|(\sum I - \langle I \rangle)|) / (\sum I)$ where $\langle I \rangle$ is the average intensity of multiple measurements.
$R_{\text{work}} = \sum_{\text{hkl}} ||F_{\text{obs}}(\text{hkl})| - F_{\text{calc}}(\text{hkl})| / \sum_{\text{hkl}} |F_{\text{obs}}(\text{hkl})|$ .
$R_{\text{free}}$ = cross validation R factor for 500 of the reflections against which the model was not refined. This represents 10% of the dataset.

### Structural analysis with Alphafold3

Using AlphaFold3 (31) we folded a single gp48 protomer with a single Fe^3+^ ion (Fig. 5F), and the predicted structure aligns well with the N-terminal 42 residues in the crystal structure. The C-terminal 30 residues are folded with lower confidence – consistent with their disorder in the structure – but the Fe^3+^ is well coordinated by residues C49, C52, C59 and H62 (Fig. 5F, S9A). We also folded gp48 as a dimer and included two Fe^3+^ ions (Fig. 5G, S9B). Each protomer of the predicted dimeric structure again aligns well with the N-terminal 42 residues mapped in the crystal structure, including the residues contributing to the dimer interface (Table S3); the RMS between the predicted and crystallographically determined structures of residues 1 – 42 per protomer is 0.37 Å (Fig. 5G). To clarify, these Fe ions are distinct from those at the plane of symmetry between two unit cells of the crystals.

Next, we predicted the dimeric structure of Mpg consistent with the dimerization of homologous proteins (29) (Fig. 6A, Table S4). The dimerization interface encompasses approximately 993.4 Å^2^ of interface area, and alignment of an Mpg protomer with the crystal structure of the homologous *Thermotoga maritima* protein shows good structural alignment (RMSD between 81 pruned atom pairs is 1.218 Å; 11.293 Å across all 236 pairs) of the active site, although the dimerization interface is somewhat distinct (Fig. 6B). We also predicted a model of two Mpg protomers, two protomers of Adephagia gp48, and two Fe^3+^ ions (Fig. 6C). Interestingly, the gp48 dimeric interaction is maintained, the dimerization interface between Mpg protomers is interrupted, and each gp48 protomer forms a robust interaction with a Mpg protomer across 5 seeds (Fig. 6C, Table S4). The C-terminal part of gp48 containing the putative Fe^3+^ ion is predicted to interact with each Mpg protomer, but the Alphafold predictions are lower confidence and its biological relevancy is questionable. Nonetheless, because Mpg is reported to be active as a dimer (29), interruption of the interaction by gp48 provides a plausible explanation for its inhibitory activity.

**Figure 6.**
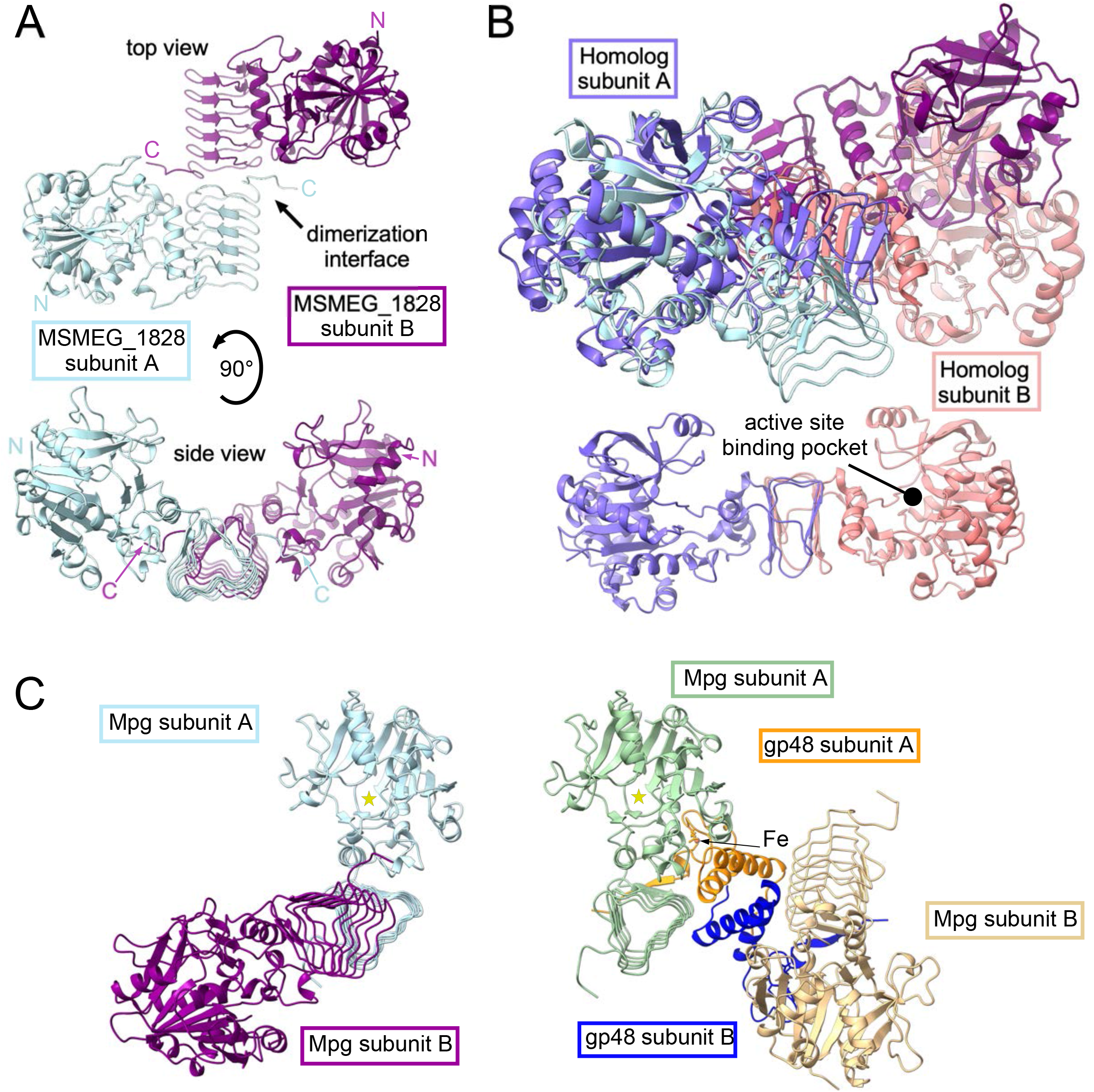
Structural insights into Msmeg_1828 and its interaction with gp48. **A.** The Alphafold3-predicted homodimer of Msmeg_1828 (Mpg) is shown from top (top) and side (bottom) views. The two subunits are colored light blue and magenta, and the N and C termini are labeled for each chain. **B.** The Alphafold3-predicted homodimer of Mpg was aligned to the crystal structure of a structural and functional homolog from *Thermotoga maritima* (PDB 2X5S) (45), which is shown in purple (subunit A) and peach (subunit B). Although the dimerization interface differs, good alignment between the enzymatic domain is observed for subunit A from both structures; the *T. maritima* enzyme active site is pointed out for the structure at the bottom of panel B. **C.** The predicted Mpg homodimer structure is shown alone (left) and in complex with an Adephagia gp48 homodimer (orange and blue) and two Fe^3+^ ions bound (right). A yellow star denotes the active site of Mpg subunit A in both structures.

### Adephagia gp48 mutants and their activities

To explore further the role of Adephagia gp48, we constructed mutant derivatives using error-prone PCR and selected for non-toxic derivatives (Fig. 7A). Avoiding large gene deletions, we sequenced 22 plasmids and identified 15 single amino acid substitutions, a nonsense mutant, four with multiple substitutions, and two C-terminal deletions (Fig. 7A, Table 3). All of the single amino acid substitutions lie within one of the N-terminal two major helices or in the short loop between them (Fig. 7A). Interestingly, we did not recover any single amino acid substitutions in the C-terminal part of the protein, suggesting it is not critical for toxicity. We confirmed the lack of toxicity of the mutants, although the A13T and E22G have less severe defects (Fig. 7B). Several C-terminal substitutions were constructed by site-directed mutagenesis (Fig. 7A), and all of these mutants retain their toxicity, showing that the gp48 C-terminus is less important for its inhibitory function (Fig. 7C).

**Figure 7.**
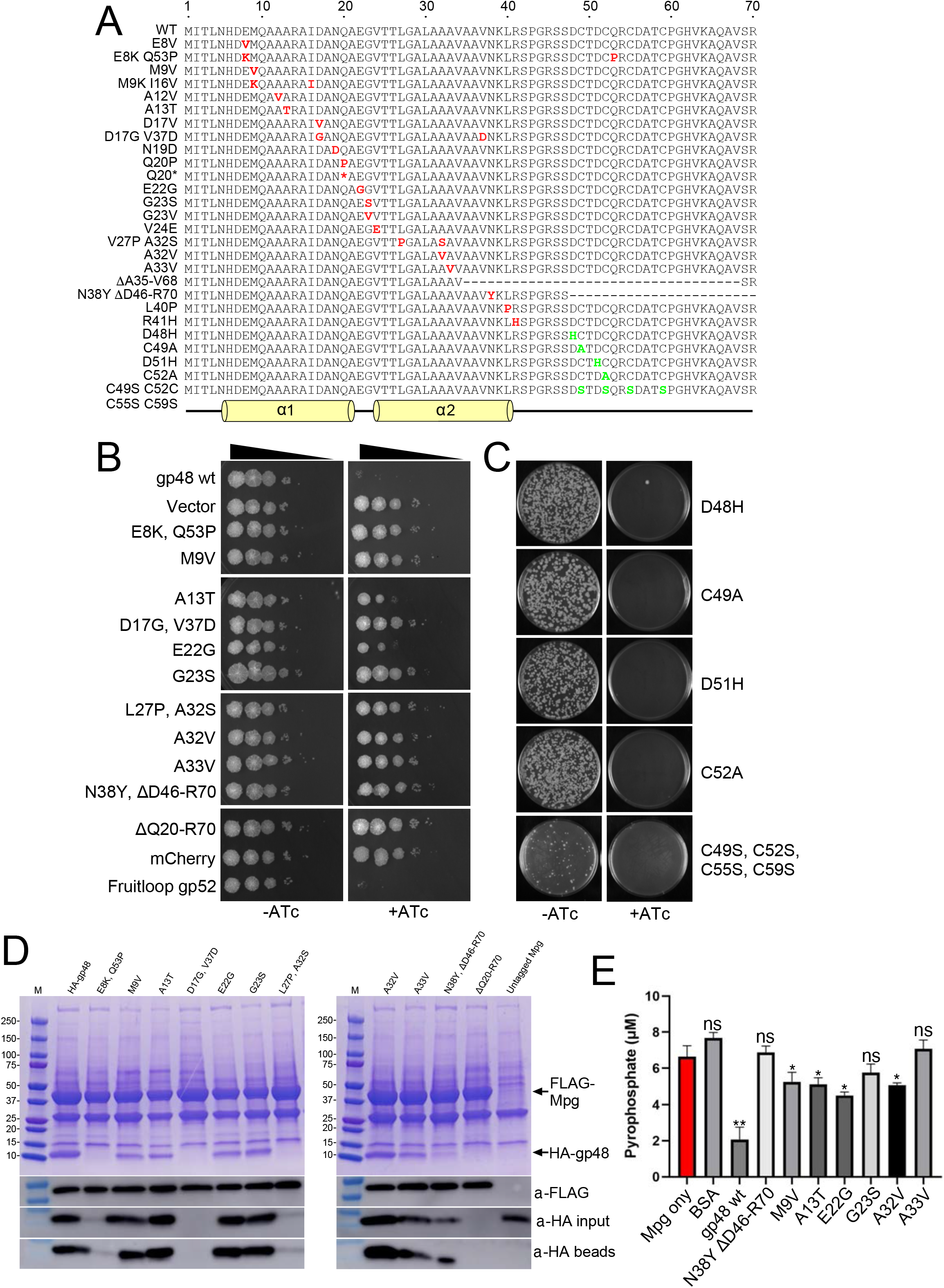
Non-toxic Adephagia gp48 mutants bind, but do not inactivate Msmeg_1828. **A.** Amino acid substitutions derived from error-prone PCR (red text) or site directed mutagenesis (green text) are shown in the amino acid sequence of Adephagia gp48. Dashes represent deleted residues and asterisk denotes a nonsense codon. **B**. Ten-fold serial dilutions of *M. smegmatis* strains expressing mutant alleles of gp48 are plated on solid media without (-ATc) or with (+ATc) inducer. **C.** Electrocompetent *M. smegmatis* mc^2^155 was transformed with plasmid carrying either single or multiple amino acid substitutions on Adephagia gp48, denoted on the right side, and an equal volume subset of recovered cells were grown with without ATc and with ATc inducer. **D**. Cell lysates expressing an HA-tagged Adephagia gp48 mutant and FLAG-tagged Msmeg_1828 (FLAG-Mpg), indicated by black arrows, were immunoprecipitated using anti-FLAG affinity columns, denatured, run on a 4-20% SDS-PAGE gel, and stained with Imperial Blue. Corresponding Western blots shown below confirm the presence of either FLAG or HA tagged protein in the input cell lysate or the anti-FLAG eluate; the primary antibody used to probe each blot is indicated on the right. **E.** Purified Adephagia gp48 (109 nM), gp48 mutants (M9V, A13T, E22G, G23S, A32V, A33V, and N38Y ΔQ46-R70 double mutant), or a BSA control (109 nM) were incubated with Msmeg_1828 (200 nM) in the presence of 500 µM GTP, 1 µM mannose-1-phosphate, and 2 mM MgCl_2_ at 25°C and pyrophosphate release was determined at 15 minutes. Statistical significance was determined by a two-tailed Student’s t-test comparing each condition to Msmeg_1828 alone (red bar), where not significant is denoted by ns; * denotes significance with *P* < 0.05.

**Table 3.** Amino acid substitutions in Adephagia gp48.

| Substitution(s) | Method<sup>1</sup> | Expression in *M. smegmatis*<sup>2</sup> | Expressed in *E. coli*<sup>3</sup> | Toxic<sup>4</sup> |
| --- | --- | --- | --- | --- |
| E8V | Error-prone PCR | NT | NT | No |
| E8K, Q53P | Error-prone PCR | No | NT | No |
| M9V | Error-prone PCR | Yes | Yes | No |
| M9V, I16V | Error-prone PCR | NT | NT | No |
| A12V | Error-prone PCR | NT | NT | No |
| A13T | Error-prone PCR | Yes | Yes | No |
| D17V | Error-prone PCR | NT | NT | No |
| D17G, V37D | Error-prone PCR | No | NT | No |
| N19D | Error-prone PCR | NT | NT | No |
| Q20P | Error-prone PCR | NT | NT | No |
| ΔQ20-R70 | Error-prone PCR | No | NT | No |
| E22G | Error-prone PCR | Yes | Yes | No |
| G23S | Error-prone PCR | Yes | Yes | No |
| G23V | Error-prone PCR | NT | NT | No |
| V24E | Error-prone PCR | NT | NT | No |
| L27P, A32S | Error-prone PCR | No | NT | No |
| A32V | Error-prone PCR | Yes | Yes | No |
| A33V | Error-prone PCR | Yes | Yes | No |
| ΔA35-V68 | Error-prone PCR | NT | NT | No |
| N38Y, ΔD46-R70 | Error-prone PCR | Yes | Yes | No |
| L40P | Error-prone PCR | NT | NT | No |
| R41H | Error-prone PCR | NT | NT | No |
| D48H | SDM | NT | NT | Yes |
| C49A | SDM | NT | NT | Yes |
| D51H | SDM | NT | NT | Yes |
| C52A | SDM | NT | NT | Yes |

<sup>1</sup>Mutations were made using either error-prone PCR or site-directed mutagenesis (SDM).
<sup>2</sup>Protein expression in *M. smegmatis* was determined by immunoprecipitation using anti-HA antibodies. NT, not tested.
<sup>3</sup>Recombinant Protein expression in *E. coli* to purify protein for biochemical pyrophosphate release assays (Fig. 7); Yes if expressed recombinantly or NT, not tested.
<sup>4</sup>Cytotoxicity was determined on *M. smegmatis* mc<sup>2</sup>155 by comparing cell viability on solid agar media with without ATc or with ATc inducer.

Several of the mutant alleles were examined for their ability to interact with Mpg *in vivo* by co-immunoprecipitation with FLAG-tagged Mpg (Fig. 7D). Most of the mutant proteins are expressed well, and are readily detectable by Western blotting (Fig. 7D). The small fragment predicted from the ΔQ20-R70 mutant was not detectable, and likely was not resolved in the gel system used. Immunoprecipitation of FLAG-Mpg pulled down all of the expressed gp48 mutants showing that all – including the N38Y ΔD46-R70 double mutant – retain their ability to interact with Mpg even though toxicity is either eliminated or greatly alleviated (Fig. 7D). Finally, we showed that all of the single amino acid substitution mutants we tested have lost the ability to strongly inhibit Mpg enzymatic activity, although the M9V, A13T, E22G and A32V mutants have milder but statistically significant reductions (Fig. 7E).

### Influence of gp48 and Mpg expression on phage infection

The potential selective advantage to the phage for gp48-mediated Mpg inactivation is unclear. To determine if gp48 expression interferes with infection by other phages we prepared a cocktail of 61 phages, infected a gp48 expressing strain of *M. smegmatis* and a control strain, isolated total RNA, and mapped the sequencing reads to each of the phages. We observed small reductions in RNA from several phages when gp48 is expressed, with the largest effect for phages Courthouse and Bxb1 (Fig. 8). We repeated this experiment, comparing the number of phage sequencing reads in a strain in which CRISPRi knocks down Mpg expression (Fig. 4E) with a control strain (Fig. 8). We observed no consistent patterns indicating inhibition of phage gene expression in either of the conditions we tested. Because gp48 expression appears to reduce the number of Bxb1 reads Bxb1 infection, we examined Bxb1 by RT-PCR (Fig. S8) but saw no difference in RNA levels. These observations suggest that either Mpg inactivation does not influence phage infection, that we failed to identify a targeted phage among those we tested, or that Mpg could be required for productive infection by one or more phages, but that it is not reflected in changes in phage gene expression.

**Figure 8.**
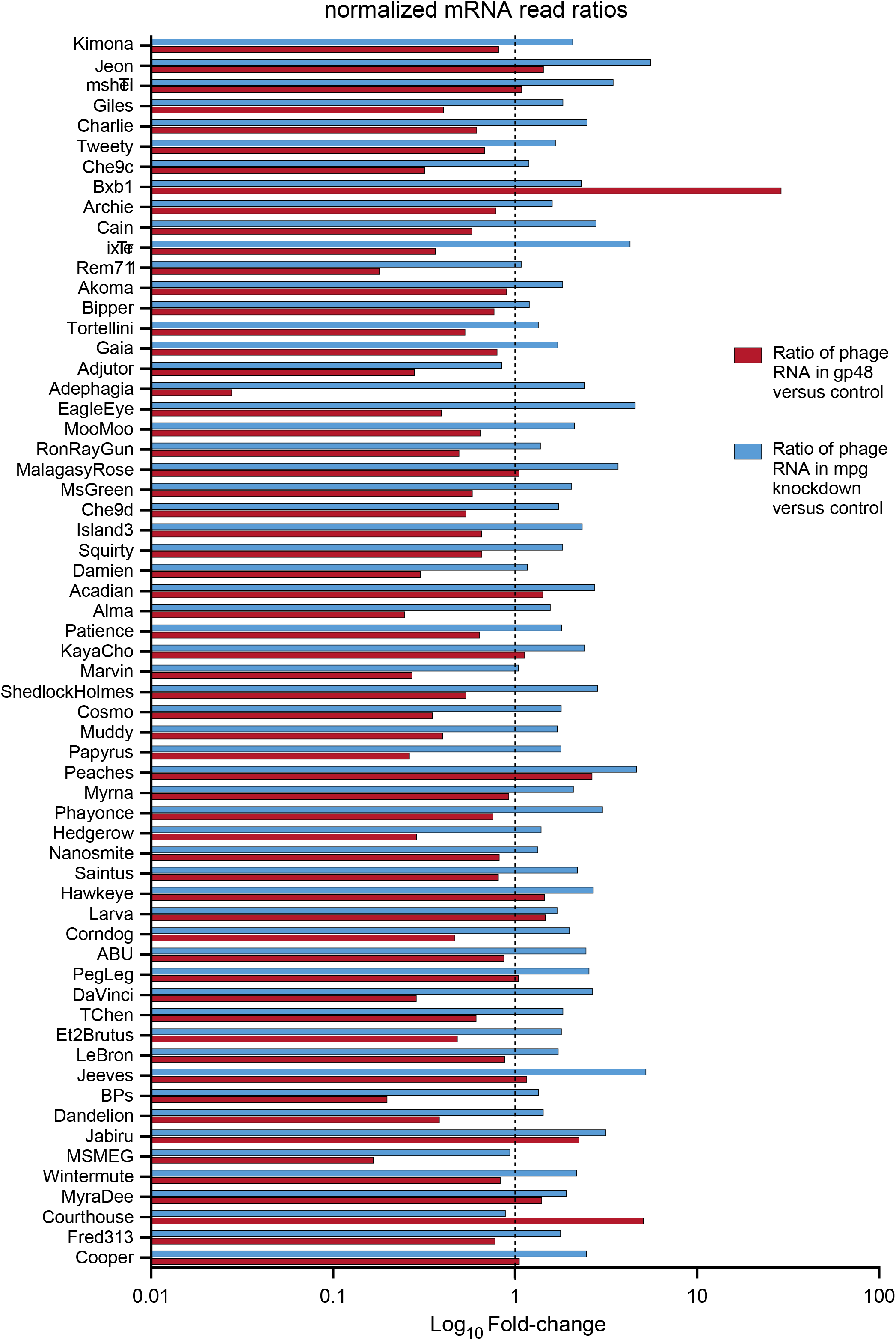
Adephagia gp48 expression does not appear to inhibit superinfection. Mid-logarithmic *M. smegmatis* cultures carrying either empty vector, expression plasmid with inducible Adephagia gp48, or a CRISPRi plasmid targeting Msmeg_1828 (Mpg) were induced with ATc for 120 minutes, then infected with a panel of 60 phages at an M.O.I ∼0.1 per phage. mRNA was extracted 150 minutes post-infection, reverse-transcribed, sequenced, and reads were aligned to each respective phage genome and normalized to the *M. smegmatis* host genome reads per sample. The ratios of normalized mRNA reads for both gp48-expressing (red) and Mpg-CRISPRi knockdown (blue) strains-to-empty vector control are plotted as log_10_ fold-changes horizontally on the x-axis for each phage, listed along the y-axis. A vertical dotted line is shown at “1” denoting a 1:1 sample-to-vector ratio of normalized mRNA reads.

## DISCUSSION

The large proportion (∼70%) of phage proteins with undefined functions presents a major challenge in understanding phage growth, dynamics, and evolution. These functionally undefined proteins are often not required for lytic growth (10, 32), suggesting that they are more likely to act in influencing phage-bacterium dynamics. Screening phage genes for toxicity provides a useful approach for functional characterization, because toxicity often arises from interaction with and inactivation of host protein functions. Here we showed that Adephagia gp48 toxicity derives from direct interaction and inactivation of *M. smegmatis* Msmeg_1828 (Mpg), interfering with GDP-mannose biosynthesis, a precursor for many cell wall constituents.

Biochemical and structural characterization of Adephagia gp48 shows activity as a dimer, with two alpha helices in the N-terminal region of each protomer engaged in dimeric interactions. The crystal structure dimer interface involves at least 24 residues in each gp48 protomer – four forming hydrogen bonds (S0 with R41, I2 with N38, and Q20 with itself) and an additional 20 buried residues with a total dimerization interface surface area of 1932 Å^2^ total – and appears quite robust, with some dimer-sized molecules observed by SDS-PAGE (Table S3). These helices, and likely the dimeric interaction itself, are needed for toxicity, as non-toxic mutants have substitutions in this protein region. The remaining C-terminal 27 residues are poorly ordered in the crystal, appear to coordinate a metal or an Fe-S cluster, but are not required for toxicity. This region also has low confidence in AlphaFold-predicted structures both alone and in complex with Mpg.

The predicted structures suggest that gp48 may destabilize dimerization of Mpg and this could plausibly lead to loss of Mpg activity. The non-toxic mutants mapping in the N-terminal region of gp48 may reduce gp48 dimerization and thus alleviate the interference of Mpg dimerization. However, these mutants still interact with Mpg, raising the possibility that gp48 still binds to Mpg, but in a way in which both Mpg and gp48 dimerization are compromised. The predicted Mpg-gp48 structure suggests that the C-terminal part of gp48 could engage in such an interaction, but this should be interpreted cautiously because the AlphaFold predictions of the gp48 C-terminal region are not of high confidence.

It is unclear what benefit inactivation of *M. smegmatis* Mpg brings to Adephagia. However, other instances of phage protein toxicity inactivate a host function that is required for phage exclusion, thus preventing other phages from competing for resources required for efficient phage replication (10, 19). The prior finding that Fruitloop gp52 prevents superinfection of phage Rosebush through inactivation of DivIVA is a good example (19). Although we couldn’t identify any phages appearing to require Mpg for infection using the transcriptomic-based assay, such phages could have been excluded from the screen. However, because GDP-Mannose is required for biosynthesis of a large array of complex glycolipids, glycopeptidolipids, and methylated polysaccharides, it seems likely that some phages would require these for infection. Recently, we’ve found that some phages require *M. smegmatis* genes implicated in synthesis of methylated polysaccharides (our unpublished observations), but if the requirement is for a late stage in infection, then it would not be detected in the RNAseq based assay. Infection of such phages could thus be compromised when Mpg is inactivated but would escape detection in an RNA-based screen. Alternatively, inactivation of Mpg could compromise the host and facilitate lysis, but we do not see any lysis defect when *48* is deleted from the Adephagia genome.

In summary, we have identified the basis of toxicity by Adephagia gp48 which binds to and inactivates an enzyme playing a central metabolic role in the mycobacteria. Adephagia gp48 also binds to and inactivates Mpg in *M. abscessus* and may act broadly in other *Mycobacterium* strains. Further exploration of phage toxic genes should reveal other bacterial targets that are essential for growth and play core roles in *Mycobacterium* physiology.

## MATERIAL AND METHODS

### Bacterial Strains and growth

*M. smegmatis* mc^2^155 strains were grown in Middlebrook 7H9 liquid medium, supplemented with 10% albumin dextrose complex (ADC) at 37° C (33). *M. abscessus* GD40 was similarly grown but supplemented with 0.0448% (v/v) oleic acid. For protein expression, Rosetta2 *E. coli* was cultured in ZYP-5052 autoinduction media [10 g/L tryptone, 5 g/L yeast extract, 1 mM MgSO_4_, 1x-trace metals (Teknova), 200-mM PO_4_, 25-mM SO_4_, 50-mM NH_4_, 100-mM Na, 50-mM K, 0.5% glycerol, 0.05% glucose, 0.2% α-lactose] supplemented with selective antibiotics as required (34).

To determine cytotoxicity, bacterial strains were grown to mid-logarithmic growth, normalized to OD_600_ ∼0.1, and ten-fold serial dilutions plated onto solid medium with or without ATc inducer.

### Plasmid constructions

Plasmids are shown in Table S1 and as described previously (17). Genes were PCR-amplified with 20 bp of 5’ and 3’ end flanking homology to vectors supplied by PCR primers and assembled into linearized vectors via Gibson Assembly, transformed into 5α *E. coli* [New England BioLabs, (NEB)] in Luria–Bertani (LB) media with selective antibiotic markers, and miniprepped using a Nucleospin plasmid purification kit (Macherey-Nagel).

### Bioinformatics

Phage genome analyses with Phamerator (23) and MMseqs2 (35) used databases Actinobacteriophage_2422 and Actino_Draftv501, respectively. Amino acid sequences were aligned using MUSCLE (36) and phylogenetic trees were constructed by maximum likelihood method (LG model 500 bootstrap replicates), and per-residue evolutionary conservation was scored using ConSurf [https://consurf.tau.ac.il/choose_params/?number=1787776374] in MSA mode.

Structural models for protein complexes were predicted by using the AlphaFold3 server (31). The highest confidence model, predicted by pLDDT scoring for monomeric models and TM-score (ipTM) for oligomeric complexes, was visualized in ChimeraX Daily (v.1.12) (37).

### Construction of AdephagiaΔ*48* and AdephagiaΔ*43*Δ*48*

AdephagiaΔ*48* and AdephagiaΔ43Δ*48* were constructed by bacteriophage recombineering of electroporated DNA (BRED). A 500 bp synthetic substrate carrying a 171 bp deletion within gene *48* and 229 bp of flanking DNA was co-electroporated with Adephagia genomic DNA into recombineering strain *M. smegmatis* (pJV138), and plated for primary plaques (24). Plaques were PCR-screened and positive plaques replated and rescreened similarly. Plaque-purified mutants were sequenced to confirm the deletion and lack of other mutations (Azenta).

### Fecundity Assay

Phage plaques were recovered as agar plugs, resuspended in 1 ml phage buffer supplemented with 1 mM CaCl_2_, vortexed, and incubated for 10 minutes at 25°C. These plaques were clarified by centrifugation, ten-fold serially diluted, and plated for plaque counts on *M. smegmatis* mc^2^155.

### Microscopy

*M. smegmatis* mc^2^155 cells carrying inducible mCherry-fusion plasmid were grown to mid-logarithmic phase, diluted to OD_600_ ∼0.1, and induced with ATc for 0 (uninduced), 2, 6, 8, 12, and 24 hours at 37°C. At each timepoint, a portion was added to SlowFade Diamond Antifade Mountant (Thermo Fisher), mounted onto a glass slide, and imaged in both brightfield and fluorescence (red channel) on a Zeiss Axiostar plus transmitted-light microscope using 100x oil-immersion objective (1000x total magnification with 10x ocular), with fluorescence illumination provided by an HBO 50 mercury arc lamp illuminator (Zeiss) with a Chromafilter set. Images were captured with an AxioCam MRc5 camera using AxioVision software (v4.6.3.0).

### Pulldown assay and identification of binding partners

*M. smegmatis* mc^2^155 or *M. abscessus* GD40 strains expressing HA- and/or FLAG-tagged proteins, or untagged controls, were induced with ATc for 4 hours at 37°C, lysed, washed twice, and eluted using a co-immunoprecipitation kit (Pierce) as previously described (12). Eluates were separated by SDS-PAGE and stained with Imperial Blue. Proteins were identified by excising gel bands, digesting with trypsin, and analyzing using nanoLC-MS/MS at the Biomedical Mass Spectrometry Center, University of Pittsburgh. Peptides were reassembled into identified proteins with MASCOT searches against a database of the Adephagia_48 protein sequence plus all relevant bacterial proteins (NCBI Reference Sequence NC_008596.1 with the corrected protein sequence of Msmeg_1828 as in Balasubramanian et al. (25) or the GD40 whole genome sequence (38)), and the data in Table 1 were statistically described with Scaffold. Tagged-proteins were identified by Western blotting as previously described (39), using either anti-HA or anti-FLAG primary antibody (Thermo Fisher) and Goat-Anti-Mouse IgG Fc (HRP) secondary antibody (Abcam), with chemiluminescence autoexposure detection on an Amersham Imager 600 (GE Healthcare).

### Purification of recombinant Adephagia gp48 and Msmeg_1828

Coding sequences for Adephagia gp48 and Msmeg_1828 were assembled into a pET28a-derived vector encoding an N-terminal 10x-histidine tag, mRuby2 fluorophore, and TEV cleavage site, and expressed in *E. coli* by autoinduction for 3 days at 17°C shaking. Cells were centrifuged for 10 minutes at 5,000 x g at 4°C, resuspended into lysis buffer with lysozyme and a protease inhibitor cocktail (Table S2), lysed by serial passage through an Emulsiflex C3 homogenizer at 17,000 psi, cleared by centrifugation for 40 minutes at 33,000 x g, and purified by Ni-NTA affinity chromatography (buffers, column volumes, and flow rates in Table S2). Excess in-house TEV protease was added to cleave the N-terminal tag, followed by a second Ni-NTA pass to remove cleaved mRuby2 and uncleaved fusion protein. Adephagia gp48 underwent a second dialysis into high-pH CAPS buffer to increase its net charge and promote binding to a subsequent anion-exchange column. Peak anion-exchange fractions (Table S2) were pooled, concentrated and buffer-exchanged using Ultra centrifugal filter devices (Amicon), and further purified by Sephacryl gel-filtration, with fractions assessed by SDS-PAGE and A280 throughout.

### Protein crystallization and structure determination

Crystals were grown at 25°C using a sitting-drop vapor diffusion technique against a well solution of 2 M ammonium sulfate, 100 mM Tris-HCl pH 7.0, and 200 mM lithium sulfate for 3 weeks, then cryoprotected in an identical buffer with elevated ammonium sulfate (4 M) prior to flash-freezing in liquid nitrogen.

High-resolution diffraction data were collected at beamline 31-IDD (Advanced Photon Source of Argonne National Laboratory) and processed using iMosflm, Phenix, RESOLVE, COOT, and MolProbity (40–44) for integration, molecular replacement (using a monomeric AlphaFold2 predicted-model as the search model), density modification, model building, and refinement validation, respectively (Table 2). Structure factors and atomic coordinates are deposited in the PDB (www.rcsb.org) under PDBid code 38LF.

### Phage interference RNA-seq

A culture of *M. smegmatis* mc^2^155 at OD_600_ ∼0.5 carrying pKF8 (vector), pKF175 (Adephagia gp48), or pML113 (sgRNA targeting *Msmeg_1828*) was induced with ATc for 120 minutes, infected with a cocktail of 60 unique phages at an M.O.I. of 0.1 per phage, and total nucleic acid was isolated 150 minutes post-infection and DNA-depleted (Thermo Fischer) and contaminating RNAs (Qiagen). A cDNA library was created by RT-PCR using random hexamers (NEB) or phage-specific capsid gene primers to confirm putative hits. Sequencing reads were aligned to a concatenated *M. smegmatis*-60-phage reference to plot strand-specific mRNA reads per phage and normalized.

### Mutagenesis of Adephagia *48*

Mutants of Adephagia gp48 were generated by error-prone PCR mutagenesis of the coding sequence using a biased dNTPs pool (5mM dATP/dGTP, 25mM dCTP/dTTP), 7 mM MgCl_2_, and OneTaq DNA Polymerase (NEB), then cloned into pKF8 via Gibson Assembly, and transformed into NEB-5α cells. The plasmid library was recovered, miniprepped, and transformed into electrocompetent *M. smegmatis* mc^2^155, which was plated on solid media with or without inducer. Colonies from induced plates PCR-screened, gel-extracted, and Sanger-sequenced to identify mutations in the *Adephagia 48* gene.

### Pyrophosphate Release Assays

Proteins were purified as in Table S2, omitting the TEV-cleavage and gel-filtration steps, then dialyzed into 50 mM Tris pH 7.5, 2 mM MgCl_2_. Pyrophosphate release was measured using a pyrophosphate assay kit (Sigma-Aldrich); reactions were incubated at 25°C and fluorescence intensity (λ_ex_/λ_em_ = 316/455 nm) was read periodically using a Synergy H1 plate reader (BioTek).

For the dose-dependent kinetic assay (Fig. 4F), 30 nM Msmeg_1828 and substrates (500 µM mannose-1-phosphate, 250 µM GTP, 2 mM MgCl_2_) were mixed with varying concentrations of purified Adephagia gp48 or a BSA control (0-1 µM), and pyrophosphate signal was measured. For the mutant assay (Fig. 7E), triplicate reactions containing purified 109 nM Adephagia gp48 mutants or BSA control were incubated with 200 nM Msmeg_1828 in the presence of 500 µM GTP, 1 µM mannose-1-phosphate, 2 mM MgCl_2_, purified Msmeg_1828 (200 nM) at 25°C, and pyrophosphate release was measured for 15 minutes. Raw reads were converted into pyrophosphate concentration by linear regression against a standard curve and fit to a nonlinear release-kinetics model in GraphPadPrism; group comparisons used a two-tailed-t-test (P <0.05 considered significant).

## ACKNOWLEDGEMENTS

We thank Jennifer Rocher, Ching-Chung Ko, Colin M. Lewis, Saeed A. Binsabaan, Daniel A. Russel, Rebecca A. Garlena, and Dorothee Liebschner for technical assistance and comments on the manuscript. This work was supported by grants from the National Institutes of Health (GM131729) and from the Howard Hughes Medical Institute (GT17580) to GFH; and from the National Institutes of Health (AI173544) to KGF.

## DATA AVAILABILITY

Structure factors and atomic coordinates for the Adephagia gp48 structure presented here can be found in the PDB (www.rcsb.org) under PDBid code 38LF.

## AUTHOR CONTRIBUTIONS

Conceptualization: MJL, KGF, GFH; Methodology: MJL, KGF; Validation: MJL, KGF, APV; Formal Analysis: MJL, KGF, APV, GFH; Investigation: MJL, KGF, J.M., DJ; Resources: APV, GFH; Data Curation: MJL, KGF; Writing – Original Draft: MJL, KGF, GFH; Writing – Review & Editing: MJL, KGF, J.M., DJ, APV, GFH; Visualization: MJL, KGF; Supervision: APV, GFH; Project Administration: MJL, KGF, GFH; Funding acquisition: APV, GFH.

