## Supplementary Materials for "Mycobacteriophage Adephagia gp48 inhibits essential host cell wall biosynthesis enzyme mannose-1-phosphate guanylyltransferase"

Running title: Adephagia gp48

Michael J. Lauer[1], Krista G. Freeman[1,2], Danny Jiang[1], Jindanuch Maneekul[1], Andrew P. VanDemark[1], Graham F. Hatfull[1,*]

[1]Department of Biological Sciences
University of Pittsburgh
Pittsburgh PA 15260

[2]Department of Physics
Case Western Reserve University
Cleveland OH 44106

**ABSTRACT**

It is common for bacteriophages to encode proteins that are strongly inhibitory to growth of the bacterial host, and about 10% of mycobacteriophage-encoded proteins have this property. Adephagia is a Cluster K1 mycobacteriophage and prior cloning and expression of 66 Adephagia non-structural genes identified 14 that are toxic when expressed in *Mycobacterium smegmatis*. One of the expressed proteins, the 70-residue gp48, is highly toxic when expressed in both *M. smegmatis* and *Mycobacterium abscessus* and acts by binding to and inactivating the function of Msmeg_1828, a mannose-1-phosphate guanylyltransferase (Mpg). Mpg is an essential enzyme for biosynthesis of GDP-mannose, a precursor of several key cell wall constituents including lipoarabinomannan and phosphatidylinositol mannosides. The crystal structure of Adephagia gp48 shows that the N-terminal 43 residues form two alpha helices that strongly promote dimer formation; the C-terminal 27 residues are disordered but are predicted to bind Fe-S clusters via cysteine and histidine residues. Adephagia gp48 inhibits Mpg guanylyltransferase activity *in vitro* and is predicted to interrupt Mpg dimer formation. The C-terminal metal binding activity of gp48 is not required for toxicity, and non-toxic mutants have substitutions in the N-terminal alpha helices that are involved in dimerization. The selective advantage of Mpg inactivation for the phage is unclear, but it may protect from competing phages that require GDP-mannose derived molecules for efficient infection.

**IMPORTANCE**

A majority of bacteriophage-encoded proteins are of unknown function. An effective strategy for gaining functional insights into these is to identify those that are toxic when expressed in the bacterial host and determine the basis of the toxicity. We show that gp48 encoded by mycobacteriophage Adephagia is toxic when overexpressed in *Mycobacterium smegmatis* and

*Mycobacterium abscessus* and the toxicity results from gp48 interaction with the host mannose-1-phosphate guanylyltransferase (Mpg) enzyme. Mpg is required for biosynthesis of GDP-Mannose, a precursor for many essential mannose-containing cellular constituents, and gp48 acts a dimer to binding to Mgp and inactivate its enzymatic activity.

**SUPPLEMENTARY MATERIALS**

**Table S1. Plasmids used in this study**

| Plasmid | Gene Insert | Replication | Expression System | Derivative |
|---|---|---|---|---|
| pKF8 | Empty Vector | Extrachromosomal | Tet-on | pTNds |
| pKF175 | Adephagia *48* | Extrachromosomal | Tet-on | pTNds |
| pKF114 | mCherry | Extrachromosomal | Tet-on | pTNds |
| pKF115 | *Fruitloop52* | Extrachromosomal | Tet-on | pTNds |
| pML50 | Adephagia *48* E8K, Q53P | Extrachromosomal | Tet-on | pTNds |
| pML51 | Adephagia *48* M9V | Extrachromosomal | Tet-on | pTNds |
| pML52 | Adephagia *48* A13T | Extrachromosomal | Tet-on | pTNds |
| pML53 | Adephagia *48* D17G, V37D | Extrachromosomal | Tet-on | pTNds |
| pML54 | Adephagia *48* E22G | Extrachromosomal | Tet-on | pTNds |
| pML55 | Adephagia *48* G23S | Extrachromosomal | Tet-on | pTNds |
| pML56 | Adephagia *48* L27P, A32S | Extrachromosomal | Tet-on | pTNds |
| pML57 | Adephagia *48* A32V | Extrachromosomal | Tet-on | pTNds |
| pML58 | Adephagia *48* A33V | Extrachromosomal | Tet-on | pTNds |
| pML59 | Adephagia *48* N38Y, ΔD46-R70 | Extrachromosomal | Tet-on | pTNds |
| pML60 | Adephagia *48* ΔQ20-R70 | Extrachromosomal | Tet-on | pTNds |
| pKF19 | HA- *Empty Vector* | Extrachromosomal | Tet-on | pTNds |
| pKF232 | HA- Adephagia *48* | Extrachromosomal | Tet-on | pTNds |
| pML61 | HA- Adephagia *48* E8K, Q53P | Extrachromosomal | Tet-on | pTNds |
| pML62 | HA- Adephagia *48* M9V | Extrachromosomal | Tet-on | pTNds |
| pML63 | HA-Adephagia *48 A13T* | Extrachromosomal | Tet-on | pTNds |
| pML64 | HA-Adephagia *48 D17G, V37D* | Extrachromosomal | Tet-on | pTNds |
| pML65 | HA-Adephagia *48 E22G* | Extrachromosomal | Tet-on | pTNds |
| pML66 | HA-Adephagia *48* G23S | Extrachromosomal | Tet-on | pTNds |
| pML67 | H*-Adephagia *48 L27P, A32S* | Extrachromosomal | Tet-on | pTNds |
| pML68 | HA-Adephagia *48 A32V* | Extrachromosomal | Tet-on | pTNds |
| pML69 | HA-Adephagia *48 A33V* | Extrachromosomal | Tet-on | pTNds |
| pML70 | HA-Adephagia *48 N38Y,* ΔD46-R70 | Extrachromosomal | Tet-on | pTNds |
| pML71 | HA- Adephagia *48* ΔQ20-R70 | Extrachromosomal | Tet-on | pTNds |
| pML4 | FLAG-Msmeg_1828 | Tweety-Integrated | Constitutive | pLO74 |
| pML5 | Empty Vector | Tweety-Integrated | Constitutive | pLO74 |
| pML6 | Msmeg_1828 | Tweety-Integrated | Constitutive | pLO74 |
| pML110 | *mCherry*-Adephagia *48* | Extrachromosomal | Tet-on | pTNds |
| pML3 | 10-His-Msmeg_1828 | Extrachromosomal | T7 (autoinduction) | pLAM3 |
| pML75 | 10-His-Adephagia *48* | Extrachromosomal | T7 (autoinduction) | pLAM3 |
| pML76 | 10-His-Adephagia *48* M9V | Extrachromosomal | T7 (autoinduction) | pLAM3 |
| pML77 | 10-His-Adephagia *48* A13T | Extrachromosomal | T7 (autoinduction) | pLAM3 |
| pML78 | 10-His-Adephagia *48* E22G | Extrachromosomal | T7 (autoinduction) | pLAM3 |
| pML79 | 10-His-Adephagia *48* G23S | Extrachromosomal | T7 (autoinduction) | pLAM3 |
| pML80 | 10-His-Adephagia *48* A32V | Extrachromosomal | T7 (autoinduction) | pLAM3 |
| pML81 | 10-His-Adephagia *48* A33V | Extrachromosomal | T7 (autoinduction) | pLAM3 |
| pML85 | 10-His-Adephagia *48 N38Y,* ΔD46-R70 | Extrachromosomal | T7 (autoinduction) | pLAM3 |
| pML111 | 10-His-mRuby2-Msmeg_1828 | Extrachromosomal | T7 (autoinduction) | pET28a |
| pML112 | 10-His-mRuby2-Adephagia *48* | Extrachromosomal | T7 (autoinduction) | pET28a |
| pML113 | sgRNA targeting *Msmeg_1828* | L5-Integrated | Tet-on | pJR962 |

**Table S2. Chromatography buffers and run parameters for Adephagia gp48 and Msmeg_1828 purification**

| Purification step | Buffer | Column/Resin/Apparatus | Adephagia gp48 | Msmeg_1828 | CV | Flow rate (ml/min) |
|---|---|---|---|---|---|---|
| Cell resuspension & lysis | Lysis | n/a | 20 mM Tris pH 8.0, 500 mM NaCl, 5% glycerol, 2 mM βME + 1mg/ml lysozyme and leupeptin, pepstatin A, aprotinin, and PMSF cocktail for resuspension | Identical | n/a | n/a |
| Affinity chromatography equilibration | Lysis | Affinity; Ni-NTA | 20 mM Tris pH 8.0, 500 mM NaCl, 5% glycerol, 2 mM βME | Identical | 10 | 10 ml/min |
| Ni-NTA wash 1 | High salt wash buffer | Affinity; Ni-NTA | 20 mM Tris pH 8.0, 1 M NaCl, 5% glycerol, 20 mM imidazole, 2 mM βME | Identical | 10 | 10 ml/min |
| Ni-NTA wash 2 | Wash buffer #1 | Affinity; Ni-NTA | 20 mM Tris pH 8.0, 200 mM NaCl, 5% glycerol, 40 mM imidazole, 2 mM βME | Identical | 5 | 10 ml/min |
| Ni-NTA wash 3 | Wash buffer #2 | Affinity; Ni-NTA | 20 mM Tris pH 8.0, 100 mM NaCl, 5% glycerol, 100 mM imidazole, 2 mM βME | Identical | 5 | 10 ml/min |
| Ni-NTA elution | Elution buffer | Affinity; Ni-NTA | 20 mM Tris pH 8.0, 100 mM NaCl, 5% glycerol, 500 mM imidazole, 2 mM βME | Identical | 10 | 0.5 ml/min |
| TEV cleavage and dialysis[1] | HEPES dialysis buffer | Dialysis; Spectra/Por 7 dialysis membrane, 3.5-kDa or 15-kDa MWCO (VWR) | 10 mM HEPES pH 8.0, 50 mM NaCl, 20 mM imidazole, 5% glycerol, 2 mM βME | 10 mM HEPES pH 8.0, 50 mM NaCl, 5% glycerol, 20 mM imidazole, 2 mM βME | n/a | n/a |
| TEV cleavage and dialysis (gp48 only)[2] | CAPS pH 11.0 dialysis buffer | Dialysis; Spectra/Por 7 dialysis membrane, 3.5-kDa MWCO (VWR) | 20 mM CAPS pH 11.0, 50 mM NaCl, 5% glycerol, 2 mM βME | N/A | n/a | n/a |
| Q column strip (pre-equilibration) | Q column buffer, 1 M NaCl | Anion-exchange; HiTrap Q HP | 20 mM CAPS pH 11.0, 1 M NaCl, 5% glycerol, 2 mM βME | 10 mM HEPES pH 8.0, 1 M NaCl, 5% glycerol, 2 mM βME | 5 | 5 ml/min |
| Q column equilibration | Q column buffer, 50 mM NaCl | Anion-exchange; HiTrap Q HP | 20 mM CAPS pH 11.0, 50 mM NaCl, 5% glycerol, 2 mM βME | 10 mM HEPES pH 8.0, 50 mM NaCl, 5% glycerol, 2 mM βME | 5 | 5 ml/min |
| Anion exchange (Q), gradient elution[3] | Q buffers, 50-1000 mM NaCl (linear gradient) | Anion-exchange; HiTrap Q HP | 20 mM CAPS pH 11.0, 50-1000 mM NaCl, 5% glycerol, 2 mM βME | 10 mM HEPES pH 8.0, 50-1000 mM NaCl, 5% glycerol, 2 mM βME | 20 | 5 ml/min |
| Protein concentration and buffer exchange | Sizing column buffer | Amicon Ultra centrifugal filter, 3-kDa MWCO (gp48) / 10-kDa MWCO (Msmeg_1828) | 10 mM HEPES pH 7.5, 200 mM NaCl, 2% glycerol, 1 mM TCEP | 10 mM HEPES pH 8.0, 50 mM NaCl, 2% glycerol, 2 mM βME | n/a | n/a |
| Size-exclusion chromatography[3] | Sizing column buffer | Gel-filtration; S-75 or S-200 | 10 mM HEPES pH 7.5, 200 mM NaCl, 2% glycerol, 1 mM TCEP | 10 mM HEPES pH 8.0, 50 mM NaCl, 2% glycerol, 1 mM TCEP | ~1 | 1 ml/min |

[1]TEV cleavage and dialysis into a buffer containing 50 mM NaCl for anion (Q) column loading.
[2]The CAPS pH 11.0 dialysis step is unique to Adephagia gp48, increasing the protein's net negative charge by raising the pH, promoting binding to the anion-exchange resin and preparing for a 50 mM NaCl anion (Q) column loading.
[3]NaCl gradient begins at ~50 mM NaCl (equivalent to 5% buffer B), matching the NaCl concentration already present in the dialysis buffer used to equilibrate the sample; CV estimated from the recorded conductivity trace (~12–33 min at 5 ml/min). Resin is HiTrap Q HP. CV = column volumes; N/D. = not determined / to be confirmed from original records.

**Table S3: Adephagia gp48 dimerization interface**

| Chain A residue[1] | Chain B residue[2] | Contact type[3] | Hydrogen bond distance (Å)[4] | Minimum inter-chain distance (Å)[2] |
|---|---|---|---|---|
| G-1 | R41 | Buried | n/a | 3.45 |
| S0 (backbone O) | R41 (side-chain $H^{\eta 12}$) | Hydrogen bond | 1.92 | 2.69 |
| V1 | N38 | Buried | n/a | 3.55 |
| I2 (backbone NH) | N38 (side-chain $O^{\delta 1}$) | Hydrogen bond | 1.92 | 2.81 |
| T3 | N38 | Buried | n/a | 5.11 |
| L4 | V34 | Buried | n/a | 3.72 |
| E8 | L30 | Buried | n/a | 4.85 |
| M9 | L30 | Buried | n/a | 3.36 |
| Q10 | L27 | Buried | n/a | 4.18 |
| A12 | L30 | Buried | n/a | 3.52 |
| A13 | T26 | Buried | n/a | 3.76 |
| I16 | T26 | Buried | n/a | 3.73 |
| D17 | T26 | Buried | n/a | 2.56 |
| Q20 (side-chain $H^{\varepsilon 22}$) | Q20 (side-chain $O^{\varepsilon 1}$) | Hydrogen bond | 2.20 | 3.02 |
| T26 | D17 | Buried | n/a | 2.56 |
| L27 | M9 | Buried | n/a | 3.58 |
| L30 | M9 | Buried | n/a | 3.36 |
| A31 | M9 | Buried | n/a | 3.97 |
| A33 | L30 | Buried | n/a | 5.00 |
| V34 | I2 | Buried | n/a | 3.68 |
| V37 | I2 | Buried | n/a | 3.55 |
| N38 (side-chain $H^{\delta 22}$) | I2 (backbone O) | Hydrogen bond | 2.04 | 2.81 |
| R41 (side-chain $H^{\eta 12}$) | S0 (backbone O) | Hydrogen bond | 1.92 | 2.69 |
| S42 | V1 | Buried | n/a | 5.29 |

[1]The chain A residue of the Adephagia gp48 homodimer determined crystallographically, defined as any residue with a heavy (non-hydrogen) atom within 5.5 Å of the opposing chain B protomer; 24 residues contribute 966 Å$^2$ per protomer to the dimerization interface using PISA crystal symmetry interface analysis; CSS = 1.000. Atoms are specified in paratheses if participating in hydrogen bonding.

[2]The closest-contacting residue on the opposing protomer (Chain A or B); distances are measured between the closest heavy-atom pair for each residue.

[3]Buried; a non-bonded contact between the residue pair. Hydrogen bond is listed if detected in the crystal structure's dimerization interface by PISA.

[4]Distance between specified atoms participating in hydrogen bonding in the dimerization interface if applicable, or n/a for not applicable.

**Table S4: Alphafold3 confidence and seed-convergence statistics Adephagia gp48 and Msmeg_1828 (Mpg) models**

| AlphaFold3 Model | Seed | Global ipTM[1] | Global pTM[2] | Ranking score[3] | Key interface[4] | Key interface ipTM[5] |
|---|---|---|---|---|---|---|
| Adephagia gp48 monomer + 1$Fe^{3+}$ (Fig. 5F) | 0 (top) | 0.86 | 0.69 | 0.89 | gp48-$Fe^{3+}$ | 0.86 |
| | 1 | 0.86 | 0.70 | 0.88 | gp48-$Fe^{3+}$ | 0.86 |
| | 2 | 0.76 | 0.58 | 0.83 | gp48-$Fe^{3+}$ | 0.76 |
| | 3 | 0.85 | 0.71 | 0.82 | gp48-$Fe^{3+}$ | 0.85 |
| | 4 | 0.83 | 0.50 | 0.78 | gp48-$Fe^{3+}$ | 0.83 |
| Adephagia gp48 dimer + 2$Fe^{3+}$ (Fig. 5G) | 0 (top) | 0.77 | 0.79 | 0.89 | itself | 0.77 |
| | 1 | 0.72 | 0.73 | 0.84 | itself | 0.73 |
| | 2 | 0.74 | 0.75 | 0.84 | itself | 0.75 |
| | 3 | 0.76 | 0.78 | 0.83 | itself | 0.77 |
| | 4 | 0.74 | 0.75 | 0.80 | itself | 0.75 |
| Mpg homodimer (Fig. 6A)[6] | 0 (top) | 0.46 | 0.66 | 0.50 | itself | 0.46 |
| | 1 | 0.46 | 0.66 | 0.50 | itself | 0.46 |
| | 2 | 0.45 | 0.65 | 0.49 | itself | 0.45 |
| | 3 | 0.43 | 0.64 | 0.47 | itself | 0.43 |
| | 4 | 0.42 | 0.64 | 0.47 | itself | 0.42 |
| Mpg dimer + gp48 dimer + 2$Fe^{3+}$ (Fig. 6C) | 0 (top) | 0.84 | 0.88 | 0.87 | gp48-Mpg (4 pairings) | 0.79-0.81 |
| | 1 | 0.80 | 0.86 | 0.84 | gp48-Mpg (4 pairings) | 0.74-0.77 |
| | 2 | 0.79 | 0.85 | 0.82 | gp48-Mpg (4 pairings) | 0.73-0.76 |
| | 3 | 0.78 | 0.85 | 0.82 | gp48-Mpg (4 pairings) | 0.73-0.76 |
| | 4 | 0.78 | 0.85 | 0.81 | gp48-Mpg (4 pairings) | 0.72-0.75 |

[1]Global ipTM; predicted interface template modeling (TM) score denoting confidence in the relative arrangement of all chains in the model (>0.8 indicates high confidence interface prediction).
[2]Global pTM, predicted TM-score, reflecting overall fold accuracy of the model.
[3]Ranking score; Composite AlphaFold3 score used to select top-ranked models from 5 independently generated seeds (0 is top-ranking, 4 is lowest-ranking).
[4]Key interface; the main interfacing protein chain with itself, another protein, or ligand in the model, depending on biological relevance.
[5]Key interface ipTM is the interface template modeling score for the biologically relevant interface as indicated in 4.
[6]PISA-derived buried interface area for the Mpg homodimer AlphaFold3 interface: 993.4 $Å^2$.

**Table S5: AlphaFold3 modeling confidence statistics for Msmeg_1828 (Mpg) dimerization of ranked 0 models**

| Per-chain confidence AlphaFold3 Model | Chain | Chain pTM[1] | Chain ipTM[2] | Max. contact prob. (n>0.5)[3] |
|---|---|---|---|---|
| Mpg homodimer (Fig. 6A-B) | Mpg (subunit A) | 0.86 | 0.46 | — |
| | Mpg (subunit B) | 0.86 | 0.46 | — |
| 2Mpg-2gp48-2$Fe^{3+}$ (Fig. 6C) | Mpg (subunit A) | 0.92 | 0.72 | — |
| | Mpg (subunit B) | 0.92 | 0.72 | — |
| | Adephagia gp48 (subunit A) | 0.64 | 0.62 | — |
| | Adephagia gp48 (subunit B) | 0.64 | 0.62 | — |
| **Pairwise interface confidence model** | **Chain interface** | **ipTM[4]** | **Min. PAE (Å)[5]** | **Max. contact prob. (n>0.5)** |
| Mpg homodimer (Fig. 6A-B) | Mpg homodimer (A-B) | 0.46 | 9.4-9.7 | 0.74 (n=48) |
| 2mpg-2gp48-2$Fe^{3+}$ (Fig. 6C) | Mpg homodimer (A-B) | 0.84 | 2.7 | 0.10 (n=0) |
| | gp48 homodimer (A-B) | 0.70 | 0.92 | — |
| | gp48(A)-mpg(A) | 0.81 | 1.5-1.9 | — |
| | gp48(A)-mpg(B) | 0.79 | 1.8-1.9 | — |
| | gp48(B)-mpg(A) | 0.79 | 1.8-1.9 | — |
| | gp48(B)-mpg(B) | 0.81 | 1.5-1.9 | — |
| | gp48(A)-$Fe^{3+}$(1), cognate | 0.53 | 4.7 | — |
| | gp48(B)-$Fe^{3+}$(2), cognate | 0.54 | 4.5 | — |
| | gp48(A)-$Fe^{3+}$(2), non-cognate | 0.29 | 8.2 | — |
| | gp48(B)-$Fe^{3+}$(1), non-cognate | 0.27 | 8.4 | — |

[1]Chain pTM; predicted interface template modeling (TM)-score per chain, demonstrating confidence in each model's own fold independent of any interface.
[2]Chain ipTM; the average interface confidence across all chains shown in the model.
[3]Max. contact prob. (n>0.5); the predicted probability that a residue pair lies within 8 Å, reported as the maximum probability observed across the interface and the number of residue pairs exceeding 0.5.
[4]ipTM; predicted interface TM-score specific to the two chains listed (>0.8 indicates high confidence interface prediction).
[5]Min. PAE; the minimum predicted alignment error between two chains; lower values indicated greater confidence in their relative position, independent of whether they are in direct contact.

### SUPPLEMENTARY FIGURE LEGENDS

**Figure S1. Pham 850 amino acid conservation and AlphaFold3-predicted tertiary structure diversity**. Primary amino acid sequences of Adephagia gp48 and five representative homologs (Illumine gp51, Amelie gp44, Hammy gp51, ZoeJ gp50, and Nergal gp46) are aligned and colored by conservation among all 144 Pham 850 members using a ConSurf evolutionary conservation mapping scheme for each amino acid, scaled 1-9; 1 is least conserved (most green), 5 is variable (white), 9 is most conserved (purple), and yellow indicates insufficient data at that aligned position. Two α-helices (α1, α2) and the C-terminal flexible region are annotated above the sequence alignment corresponding to the AlphaFold3-predicted monomeric structure of and primary sequence of Adephagia gp48. Below, monomeric AlphaFold3 models of each selected representative member of Pham 850 (Adephagia g48, Illumine gp51, Amelie gp44, Hammy gp51, ZoeJ gp50, and Nergal gp46) are shown for reach structure with residue conservation color-mapped as in the Adephagia gp48 reference scheme above.

**Figure S2. Whole Pham 850 sequence alignment.** Primary amino acid sequences from all 144 Pham 850 members were aligned in Jalview by pairwise identity, with Adephagia gp48 (red star) shown at the top as the reference sequence and residues are colored by precent identity (purple = 100%, white = 0%). Two α-helices (α1, α2) and the C-terminal flexible region are annotated above the alignment using Adephagia gp48 residue numbering, corresponding to the same predicted domains as shown in Figure S1. Conservation, alignment quality, and consensus are plotted for all 144 sequences below the alignments where sequences ranged from 63 to 93 residues (mean = 72.78 residues), standard deviation = 7.35, median = 70.

**Figure S3. Adephagia gp48 is not involved in prophage-mediated defense.** Ten-fold serial dilutions from a diverse panel of 57 unique phages spanning unrelated clusters were spotted on lawns of *M. smegmatis* $mc^2155$ lysogens carrying wildtype Adephagia (leftward panel) or Δ48 (middle panel) and incubated at 37°C for 3 days. For a focused subset of 20 of these phages, a wildtype *M. smegmatis* $mc^2155$ control lawn is indicated on the right. Phages are listed by row with their cluster and subcluster designation in parentheses.

**Figure S4. mCherry-Adephagia gp48 fusion expression retains cytotoxic to *M. smegmatis*.** Electrocompetent *M. smegmatis* $mc^2155$ cells were transformed with 200 ng plasmid carrying a construct of Adephagia *48* coding sequence with an mCherry fusion tag on the N-terminus of the expressed protein, creating pML110. Recoveries of 100 µl cells under hygromycin B selection are shown on either -ATc or +ATc solid agar media to the left or right respectively.

**Figure S5. Identity of *M. smegmatis* and *M. abscessus* GD40 co-immunoprecipitations. A.** Clarified lysates were prepared from *M. smegmatis*, expressing either untagged Adephagia gp48 (-) or Adephagia gp48 with an N-terminal HA tag (+). Following immunoprecipitation and washing, proteins were separated on a 4-20% SDS-PAGE gel and stained with Imperial Blue. An arrow indicates the expected migration of Adephagia HA-gp48, confirmed by anti-HA primary antibody in the left-hand Western blot; scissors indicate where the membrane was physically cut after transfer to separate the anti-FLAG-probed (top, unrelated samples) and anti-HA-probed (bottom) portions. **B.** Lysates of N-terminally HA-tagged gp48 were expressed with FLAG-tagged (+) or untagged (-) Msmeg_1828 and immunoprecipitated with anti-FLAG beads, washed, eluted, and separated by SDS-PAGE, and stained. An arrow indicates the presence of HA-gp48 and FLAG-Msmeg_1828. A Western blot using anti-FLAG primary antibody confirms FLAG-Msmeg_1828 identity; HA-gp48 identity was instead confirmed by LC-MS/MS of the corresponding 10-kDa gel band, with 43.0% sequence coverage (Table 1). **C.** Clarified lysates of *M. abscessus* GD40 expressing either untagged gp48 (-) or HA-gp48 were immunoprecipitated similarly to panels A and B. An arrow indicates the expected migration of Adephagia HA-gp48, which is confirmed using anti-HA primary antibody in the left-hand Western blot with scissors to mark a physical membrane separation after transfer.

**Figure S6. Adephagia C-terminus spectral scans. A.** Primary amino acid residues of Adephagia gp48 are aligned and colored by conservation among all 144 Pham 850 members using a ConSurf evolutionary conservation mapping skin for each amino acid, scaled 1-9; 1 is least conserved (most green), 5 is variable (white), 9 is most conserved (purple), and yellow is variable. A c-terminal “CTDCQRCDATC” motif containing four cystine residues is enlarged from residues 49 to 59. **B.** Ribbon diagram of the monomeric AlphaFold3 Adephagia gp48 model with selected residues C49, C52, C59, and H62 shown as sticks, and residue conservation color-mapped onto the surface, colored by the ConSurf conservation scheme as in the alignments above. **C.** Spectral scan plotting normalized absorbance units from 200 to 550 nm of purified Adephagia gp48 protein from various Q-column and S-75 fractions. A broad shoulder peak can be observed in Q fraction 9 (colored blue) from 350-500 nm, while a peak is absent for concentrated S-75 fractions 24-34 (colored red).

**Figure S7. Affinity and anion-exchange protein purification of recombinant Msmeg_1828 from *E. coli*. A.** SDS-PAGE of Msmeg_1828 (Mpg) purification from E. coli using Ni-NTA affinity chromatography. Lanes were loaded as: Whole-cell lysate (L), soluble lysate fraction (S), lysate wash (LW), high salt wash (HS), imidazole wash (W1), elution (E), TEV protease-cleaved 10x-His-mRuby2-Msmeg_1828 fusion protein (mRuby-Mpg), and cleaved Mpg are indicated above the gel. **B.** Subsequent anion-exchange (Q column) purification of TEV-cleaved Mpg: Left; chromatogram UV absorbance at OD280 (blue) with conductivity (red) over a steeply increasing NaCl gradient and peak collected fractions 1-24 indicated in between dotted lines. Right; SDS-PAGE of input fractions, flowthroughs ($F_1$, $F_2$), and corresponding peak Q-column elution fractions 1-24. **C.** SDS-PAGE of pooled and concentrated Q-column fractions (1–24) from the post-TEV cleavage sample of mRuby2-Mpg expressed from the pML111 plasmid).

**Figure S8. Expression of Adephagia gp48 in *M. smegmatis* does not defend against phage infection.** Real-time (RT) PCR validation of RNA-seq results (Fig. 8) using primers specific to the Bxb1 gene *14* (major capsid protein gene) transcript or *M. smegmatis dnaA* gene as a host control, performed on cells cDNA template from *M. smegmatis* cultures carrying empty vector (pKF8) or an inducible Adephagia gp48 expression plasmid (pKF175), infected with candidate phage Bxb1. All reactions included reverse transcriptase (RT); a single ~300 bp product was amplified for both the Bxb1 gene *14* and *dnaA* primer sets, regardless of gp48 expression, confirming successful RT-PCR detection of both phage and host transcripts, but determining that gp48 expression does not measurably alter Bxb1 transcript levels.

**Figure S9. Predicted aligned error (PAE) for AlphaFold models of Adephagia gp48 and Msmeg_1828 (Mpg). A.** PAE for Adephagia gp48 monomer + 1$Fe^{3+}$ (as shown in Fig. 5F). **B.** PAE for Adephagia gp48 dimer + 2$Fe^{3+}$ (Fig. 5G). **C.** PAE for the Msmeg_1828 (Mpg) homodimer alone (Fig. 6A). D. PAE for the full Msmeg_1828 (Mpg) dimer + Adephagia gp48 dimer + 2$Fe^{3+}$ (Fig. 6C). Global confidence metrics (ipTM, pTM, ranking score) are shown in each panel; A and B additionally report mean per-residue pLDDT for the N-terminal (1-43) and C-terminal (44-70) regions of gp48 and C and D report the Mpg-Mpg contact probability. Darker blue indicates lower predicted error and higher confidence; all panels are scaled identically (0-30 Å) to allow direct comparison.

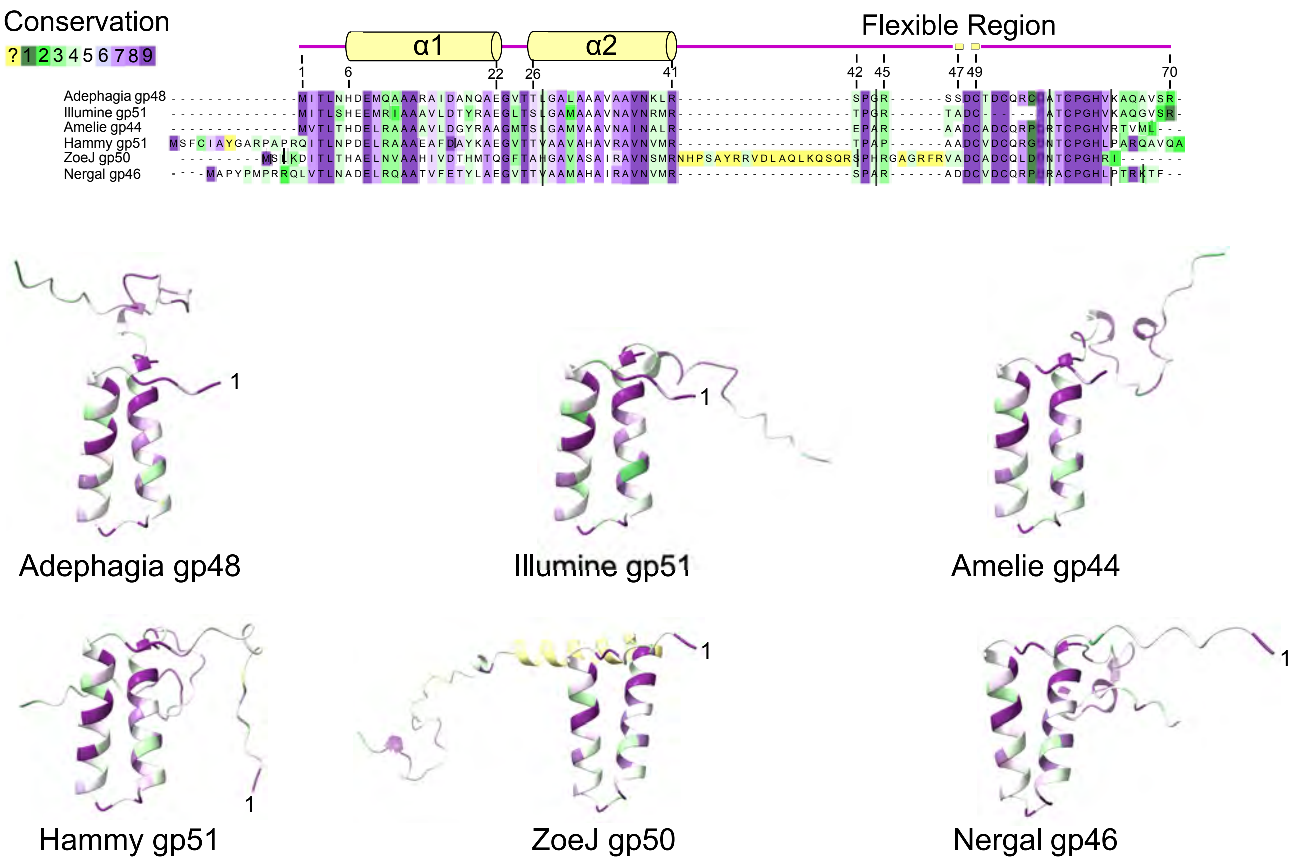
Conservation
? 1 2 3 4 5 6 7 8 9
α1
α2
Flexible Region
1
6
22
26
41
42
45
47
49
70
Adephagia gp48
Illumine gp51
Amelie gp44
Hammy gp51
ZoeJ gp50
Nergal gp46
1
Adephagia gp48
Illumine gp51
Amelie gp44
Hammy gp51
ZoeJ gp50
Nergal gp46

Figure S1

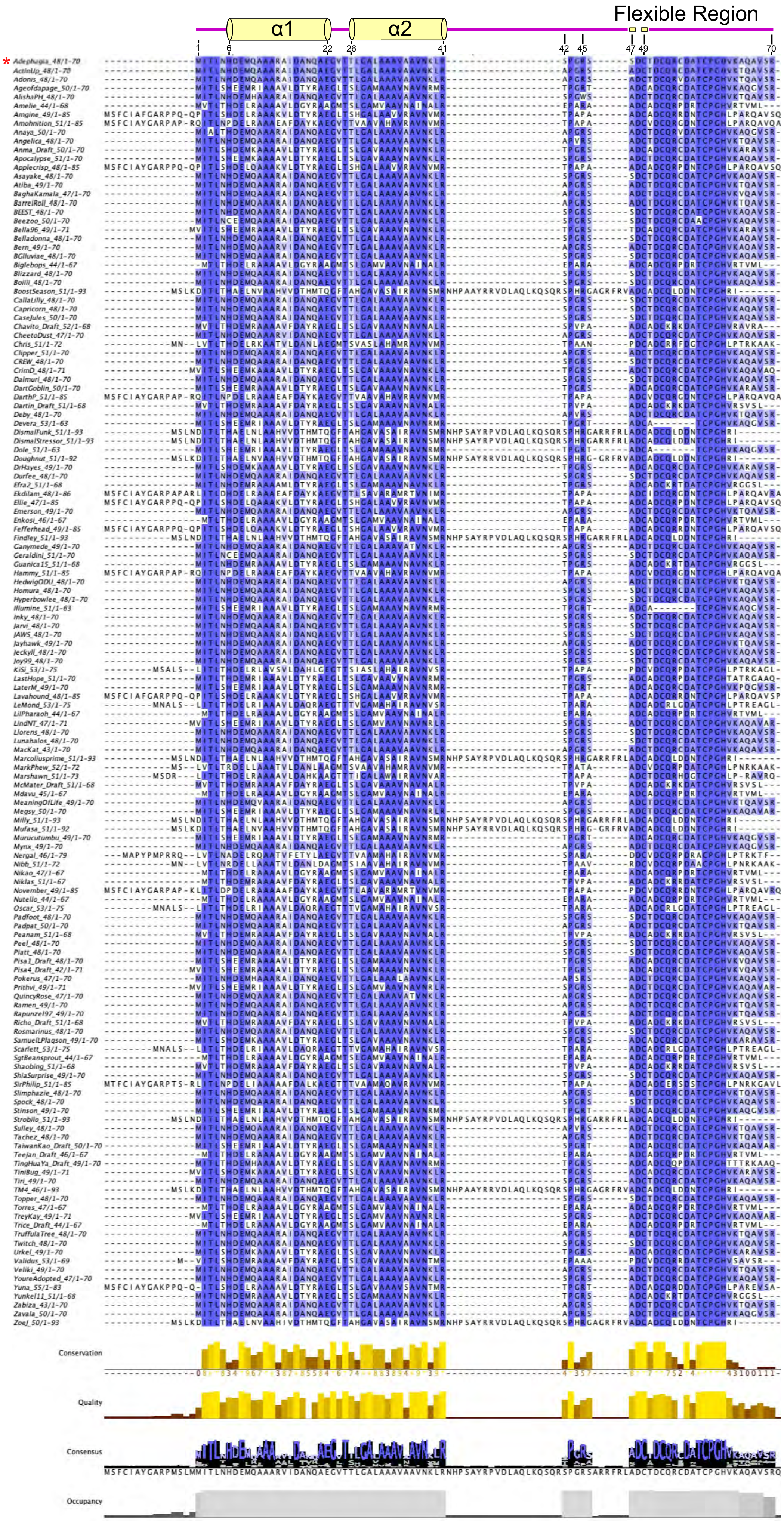
Flexible Region
α1
α2
1
6
22
26
41
42
45
47
49
70
Conservation
Quality
Consensus
Occupancy

Figure S2

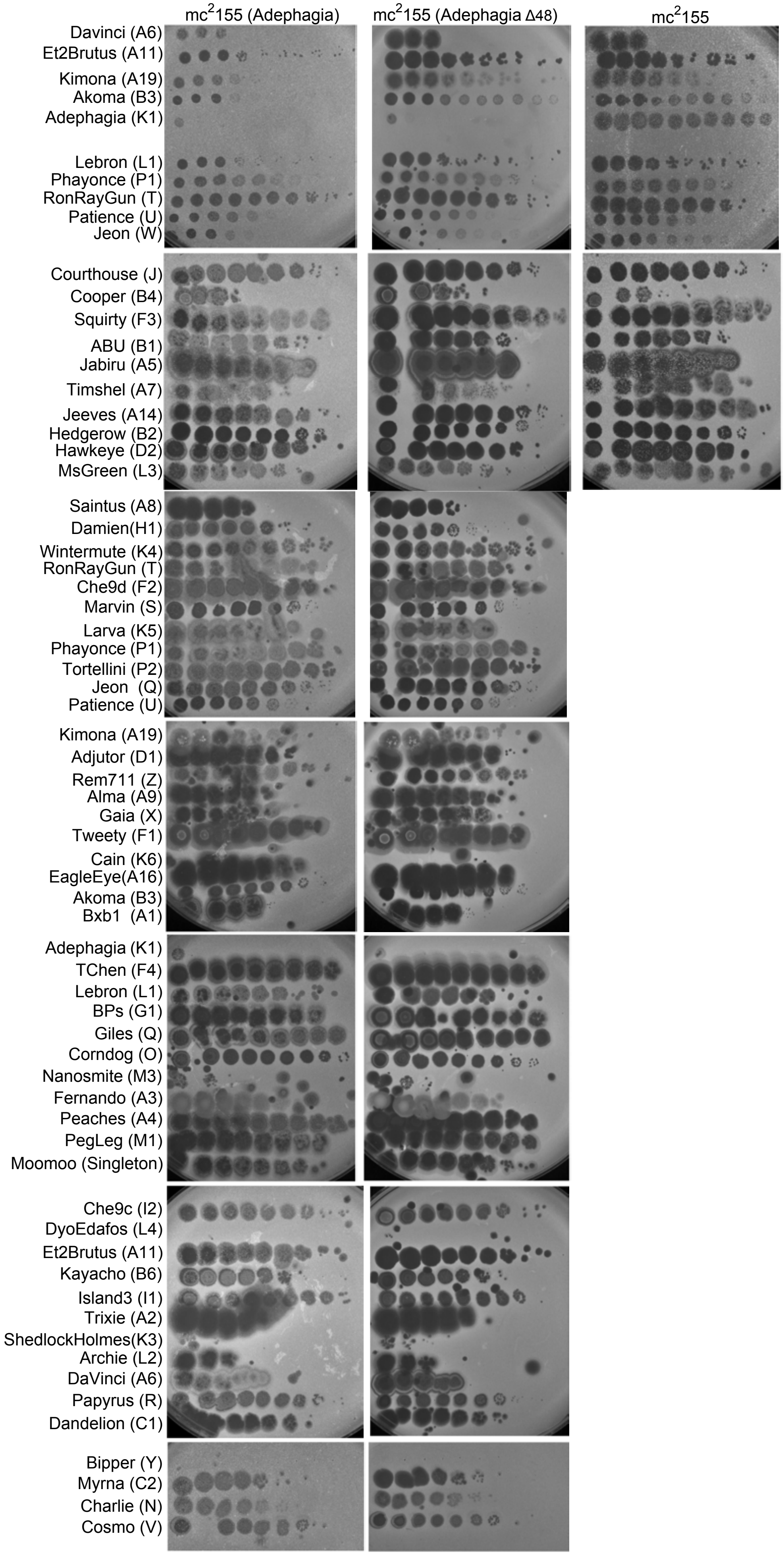
mc2155 (Adephagia)
mc2155 (Adephagia Δ48)
mc2155
Davinci (A6)
Et2Brutus (A11)
Kimona (A19)
Akoma (B3)
Adephagia (K1)
Lebron (L1)
Phayonce (P1)
RonRayGun (T)
Patience (U)
Jeon (W)
Courthouse (J)
Cooper (B4)
Squirty (F3)
ABU (B1)
Jabiru (A5)
Timshel (A7)
Jeeves (A14)
Hedgerow (B2)
Hawkeye (D2)
MsGreen (L3)
Saintus (A8)
Damien(H1)
Wintermute (K4)
RonRayGun (T)
Che9d (F2)
Marvin (S)
Larva (K5)
Phayonce (P1)
Tortellini (P2)
Jeon (Q)
Patience (U)
Kimona (A19)
Adjutor (D1)
Rem711 (Z)
Alma (A9)
Gaia (X)
Tweety (F1)
Cain (K6)
EagleEye(A16)
Akoma (B3)
Bxb1 (A1)
Adephagia (K1)
TChen (F4)
Lebron (L1)
BPs (G1)
Giles (Q)
Corndog (O)
Nanosmite (M3)
Fernando (A3)
Peaches (A4)
PegLeg (M1)
Moomoo (Singleton)
Che9c (I2)
DyoEdafos (L4)
Et2Brutus (A11)
Kayacho (B6)
Island3 (I1)
Trixie (A2)
ShedlockHolmes(K3)
Archie (L2)
DaVinci (A6)
Papyrus (R)
Dandelion (C1)
Bipper (Y)
Myrna (C2)
Charlie (N)
Cosmo (V)

Figure S3

pML110 (mCherry-gp48)

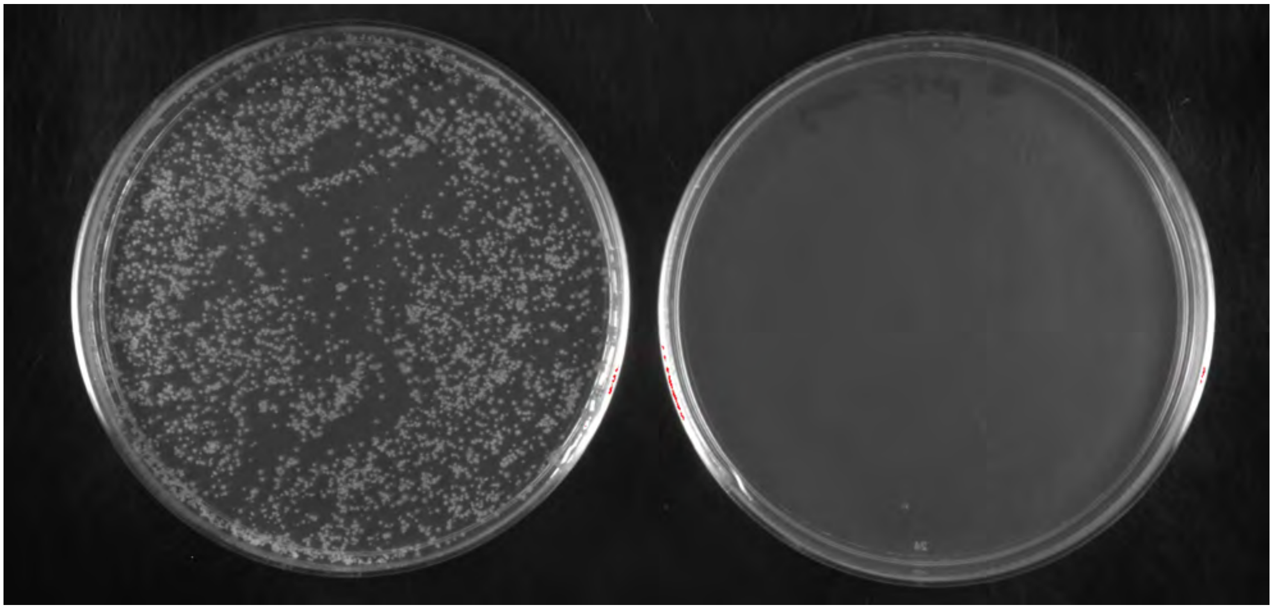

-ATc +ATc

Figure S4

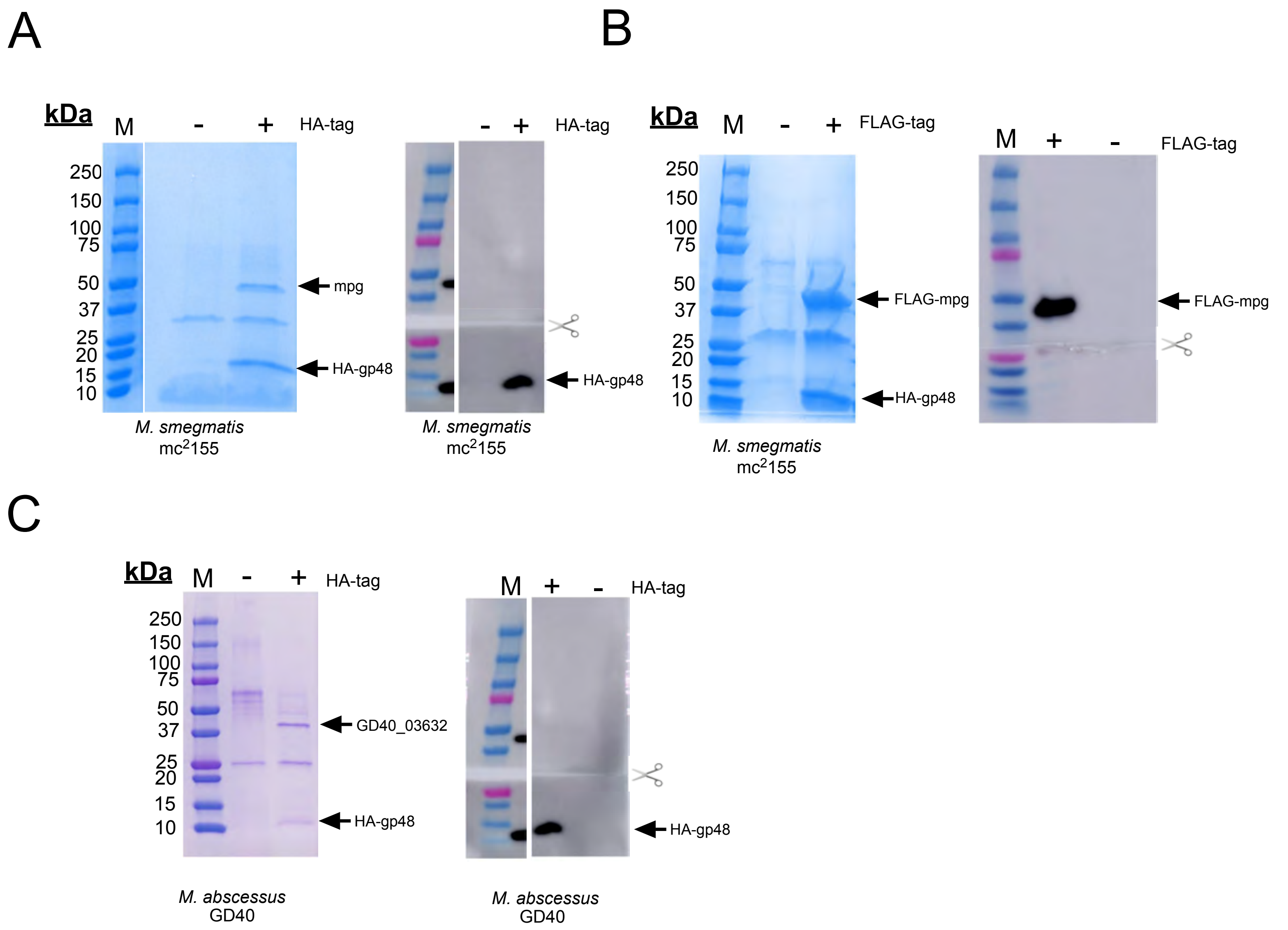
A
kDa
M
-
+
HA-tag
250
150
100
75
50
37
25
20
15
10
mpg
HA-gp48
M. smegmatis
mc2155
-
+
HA-tag
HA-gp48
M. smegmatis
mc2155
B
kDa
M
-
+
FLAG-tag
250
150
100
75
50
37
25
20
15
10
FLAG-mpg
HA-gp48
M. smegmatis
mc2155
M
+
-
FLAG-tag
FLAG-mpg
C
kDa
M
-
+
HA-tag
250
150
100
75
50
37
25
20
15
10
GD40_03632
HA-gp48
M. abscessus
GD40
M
+
-
HA-tag
HA-gp48
M. abscessus
GD40

Figure S5

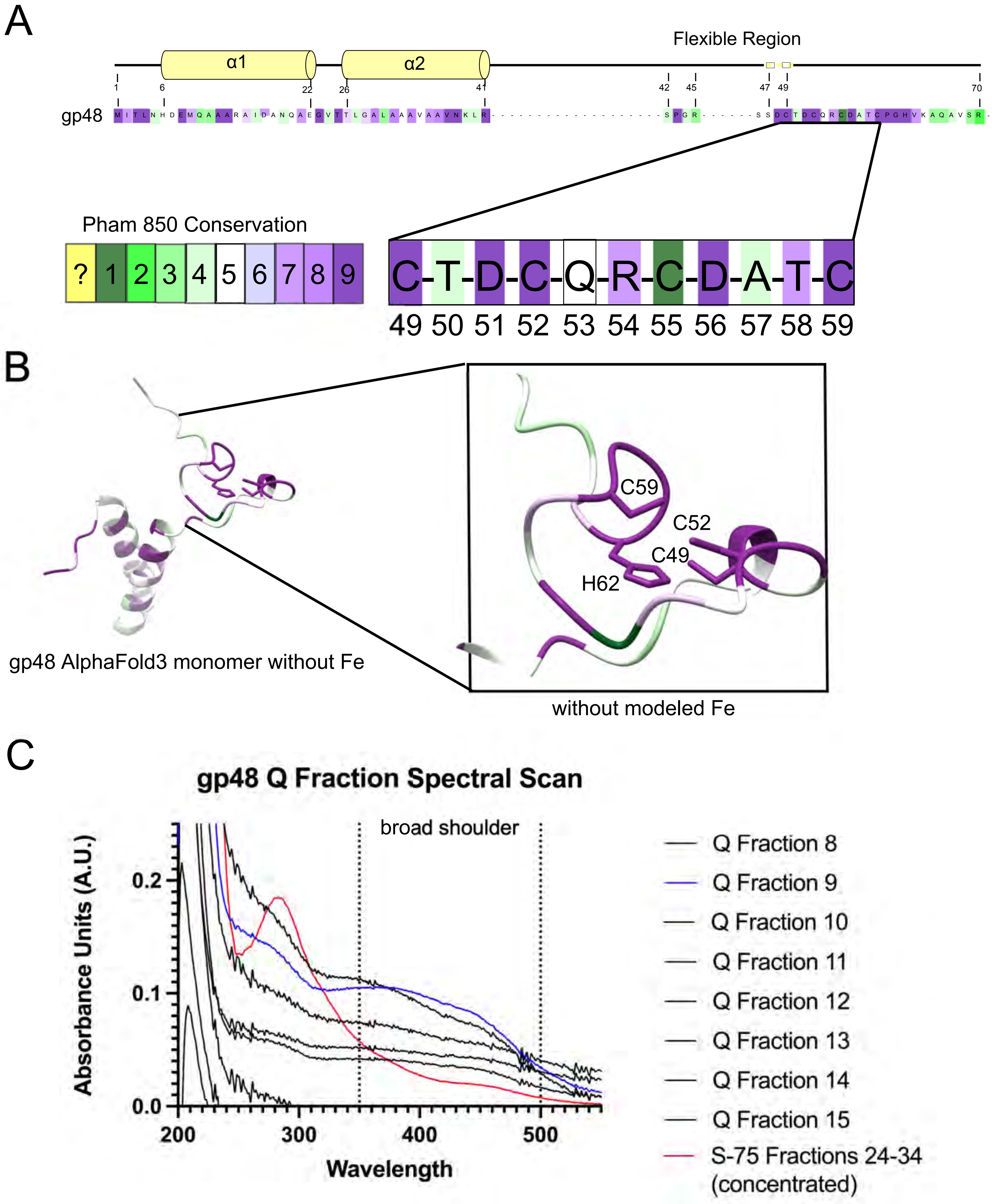
A
Flexible Region
α1
α2
1
6
22
26
41
42
45
47
49
70
gp48
Pham 850 Conservation
? 1 2 3 4 5 6 7 8 9
C-T-D-C-Q-R-C-D-A-T-C
49 50 51 52 53 54 55 56 57 58 59
B
C59
C52
C49
H62
gp48 AlphaFold3 monomer without Fe
without modeled Fe
C
gp48 Q Fraction Spectral Scan
broad shoulder
Absorbance Units (A.U.)
0.0
0.1
0.2
200
300
400
500
Wavelength
Q Fraction 8
Q Fraction 9
Q Fraction 10
Q Fraction 11
Q Fraction 12
Q Fraction 13
Q Fraction 14
Q Fraction 15
S-75 Fractions 24-34 (concentrated)

Figure S6

A

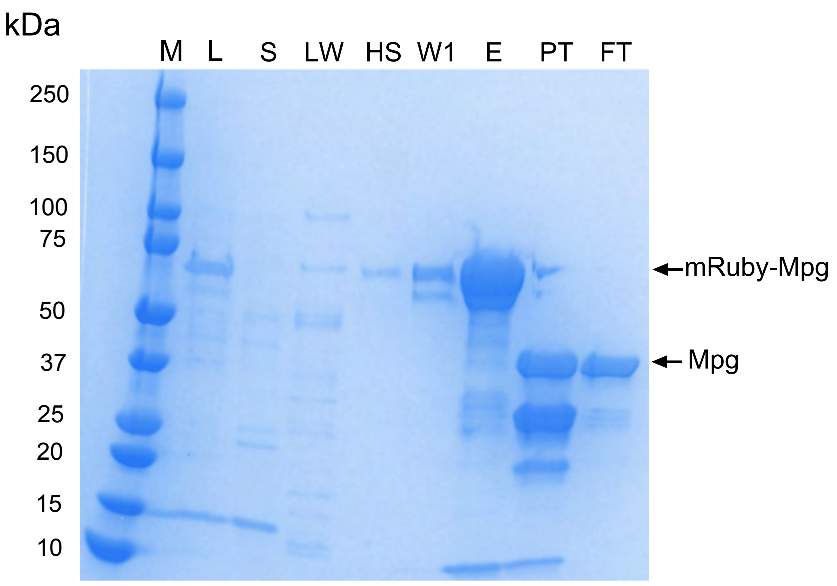


B

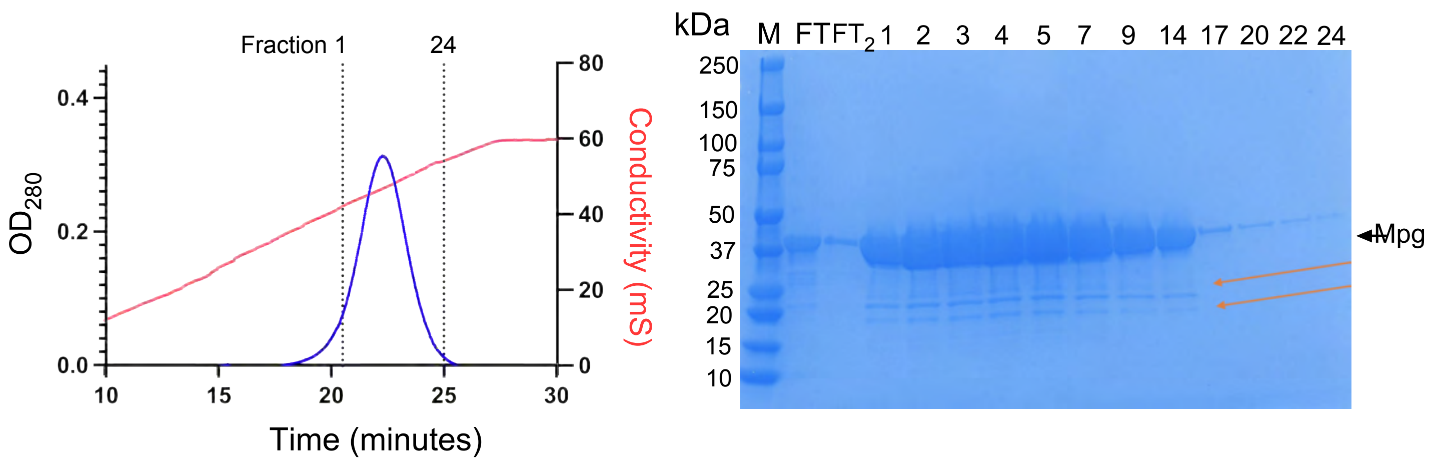


C

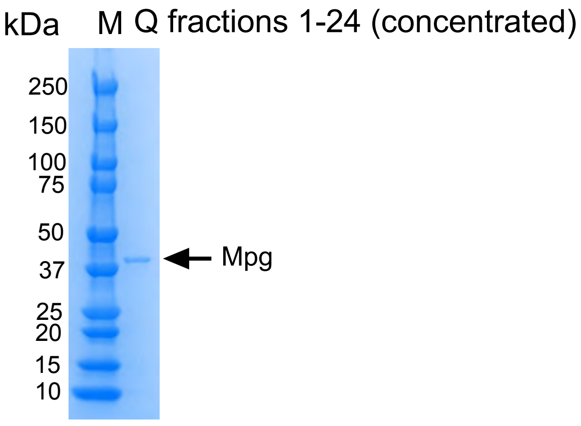


E. coli post-mRuby cleavage with TEV (pML111)

Figure S7

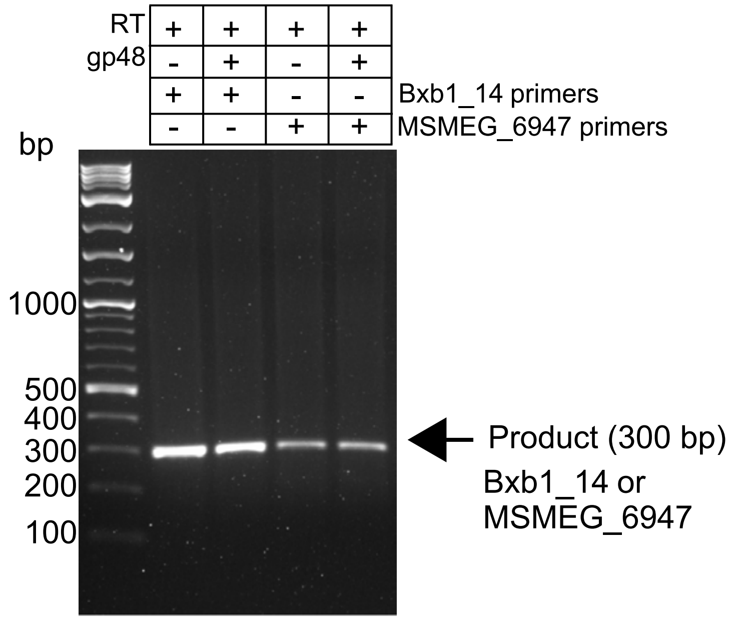
RT
+ + + +
gp48
- + - +
+ + - -
Bxb1_14 primers
- - + +
MSMEG_6947 primers
bp
1000
500
400
300
200
100
Product (300 bp)
Bxb1_14 or
MSMEG_6947

Figure S8

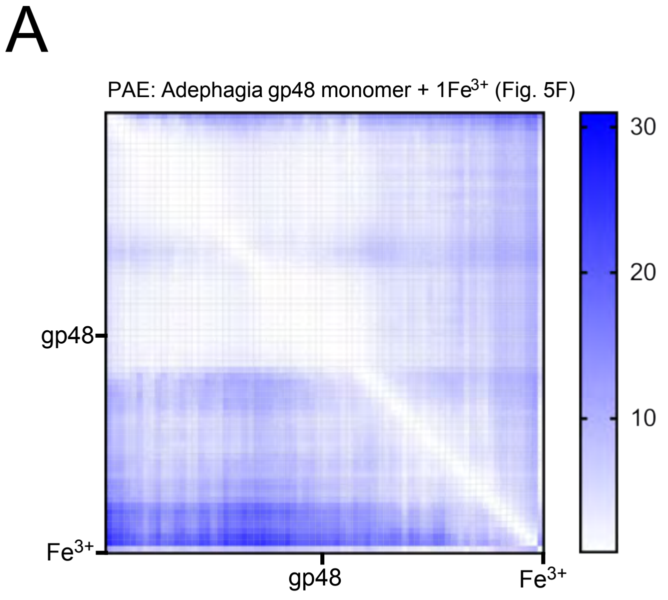


ipTM = 0.86 | pTM = 0.70 | ranking score = 0.89
Mean pLDDT: N-term (1–43) = 89.4, C-term (44–70) = 68.7

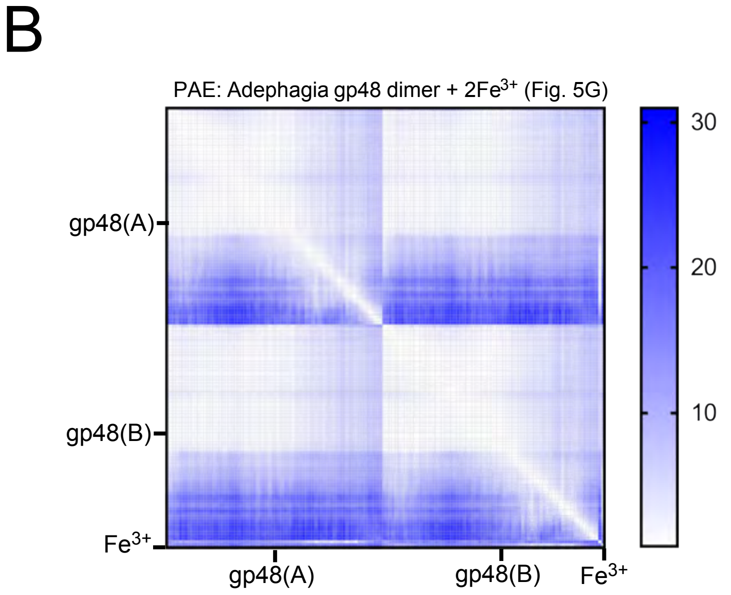


ipTM = 0.77 | pTM = 0.79 | ranking score = 0.89
Mean pLDDT: N-term (1–43) = 94.5, C-term (44–70) = 72.0

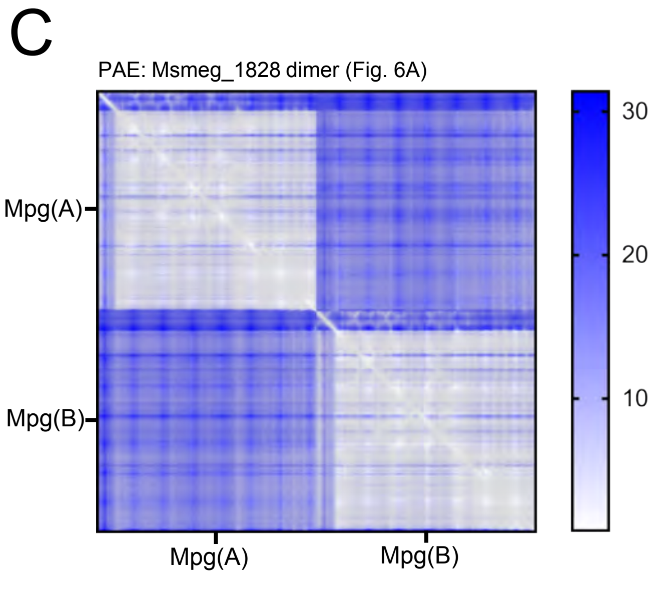


ipTM = 0.46 | pTM = 0.66 | ranking score = 0.50
Mpg–Mpg contact probability: 0.74 (48 residue pairs > 0.5)

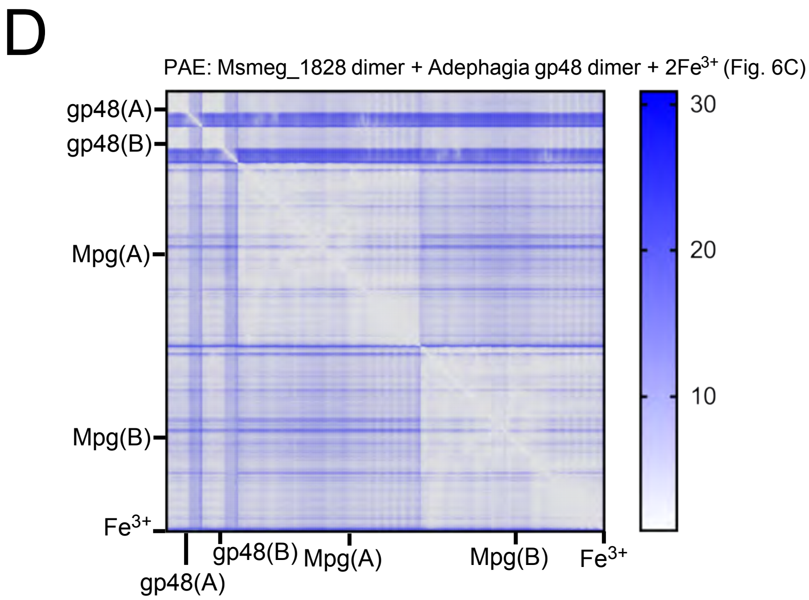


ipTM = 0.84 | pTM = 0.88 | ranking score = 0.87 Mpg–Mpg contact probability: 0.10 (0 residue pairs > 0.5)
gp48–Mpg interfaces: ipTM = 0.79–0.81 (all four pairings)

Supplemental Figure 9
